# Rem2 regulates distinct homeostatic mechanisms in visual circuit plasticity

**DOI:** 10.1101/208975

**Authors:** Anna R. Moore, Sarah E. Richards, Katelyn Kenny, Leandro de Oliveira Royer, Urann Chan, Kelly Flavahan, Stephen D. Van Hooser, Suzanne Paradis

## Abstract

Activity-regulated genes sculpt neural circuits in response to sensory experience. These calcium-sensitive genes generally fall into two categories: transcription factors and proteins that function at synapses. Yet little is known about activity-regulated, cytosolic proteins that transduce signals between the neuronal membrane and the nucleus. Using the visual system as a model, we investigated the role of the activity-regulated, non-canonical Ras-like GTPase Rem2 *in vivo*. We demonstrate that *Rem2*^-/-^ mice fail to exhibit normal ocular dominance plasticity during the critical period. At the circuit level, cortical layer 2/3 neurons in *Rem2*^-/-^ mice show deficits in both postsynaptic scaling up of excitatory synapses and misregulation of intrinsic excitability. Further, we reveal that Rem2 plays a novel, cell-autonomous role in regulating neuronal intrinsic excitability. Thus, Rem2 is a critical regulator of neural circuit function and distinct homeostatic plasticity mechanisms *in vivo*.

**HIGHLIGHTS:**

- Rem2 is required in excitatory cortical neurons for normal ocular dominance plasticity
- Rem2 regulates postsynaptic homoeostatic synaptic scaling up
- Rem2 alters the intrinsic excitability of neurons in a cell-autonomous manner

## INTRODUCTION

Sensory experience plays a key role in shaping neuronal circuitry by affecting the synaptic connectivity and intrinsic properties of individual neurons in response to sensory stimuli (Katz and Shatz, 1996). Plasticity of ocular dominance (OD) in the mammalian visual system, induced by monocular eyelid suture, is one of the best-studied models of the influence of experience on neural circuit development. The current consensus view holds that OD plasticity is achieved through an “early-phase” decrease in responsiveness to the deprived, contralateral eye through Hebbian-like LTD mechanisms (Cooke and Bear, 2010; Crozier et al., 2007; Kirkwood et al., 1996; Rittenhouse et al., 1999; Yoon et al., 2009), followed by a “late-phase” homeostatically-regulated increase in responsiveness to the open, ipsilateral eye through synaptic scaling up and increases in intrinsic excitability (Kaneko et al., 2008b; Lambo and Turrigiano, 2013; Mrsic-Flogel et al., 2007; Smith et al., 2009). Further, changes in gene expression underlie OD plasticity and the long-lasting nature of both Hebbian and homeostatic plasticity (Turrigiano, 2008). However, the molecular programs that underlie specific mechanisms of experience-dependent plasticity remain poorly understood.

Homeostatic mechanisms work to stabilize neuronal and circuit function around a set level of activity in response to perturbation (Turrigiano et al., 1998). The most extensively studied form of homeostatic plasticity is the regulation of postsynaptic strength in excitatory synapses. Postsynaptic excitatory responses can either be strengthened (scaled up) or weakened (scaled down) in response to altered network activity to maintain the overall firing rate of neurons (Desai et al., 2002; Maffei and Turrigiano, 2008; Turrigiano et al., 1998). In conjunction with regulating synaptic strengths, neurons also respond to decreased activity by modulating their ionic conductances in order to re-establish their appropriate firing rate (Aizenman et al., 2003; Desai et al., 1999; Liu et al., 1998; Marder and Prinz, 2003; Pratt and Aizenman, 2007; Siegel et al., 1994). These two homeostatic responses (i.e. regulation of intrinsic excitability and synaptic scaling) may work cooperatively to boost responsiveness to both the deprived and non-deprived eyes following monocular deprivation (Lambo and Turrigiano, 2013; Maffei and Turrigiano, 2008; Nataraj et al., 2010). Whether or how distinct signaling mechanisms regulate these individual homeostatic processes to instruct circuit function remains to be determined.

Homeostatic plasticity mechanisms are triggered by changes in intracellular calcium levels, which reflect variations in the activity level of individual neurons in a circuit (Turrigiano, 2008). Consistent with this idea, a number of molecules that have been implicated in homeostatic synaptic scaling (e.g. Arc, TNFα, MHCI, and CaMKIV) are also regulated by neuronal activity either at the level of mRNA expression, protein function or both (reviewed in Fernandes and Carvalho, 2016; Pozo and Goda, 2010). Despite the identification of these genes as regulators of synaptic scaling, a comparable molecular description of genes that regulate homeostatic changes in intrinsic excitability is lacking.

The activity-regulated Ras-like GTPase Rem2 is an excellent candidate to link the activity of a neural circuit with structural and functional plasticity. Rem2 is a member of the Rad/Rem/Rem2/Gem/Kir (RGK) protein family, a Ras-related subfamily of small GTPases (Finlin et al., 2000) whose expression and function are regulated by neuronal activity in response to calcium influx through voltage-gated calcium channels (Ghiretti et al., 2014). Further, gene knockdown approaches in cultured cortical neurons and *Xenopus laevis* optic tectum established that Rem2 is a positive regulator of synapse formation and a negative regulator of dendritic complexity (Ghiretti et al., 2013; Ghiretti et al., 2014; Ghiretti and Paradis, 2011; Moore et al., 2013), suggesting Rem2 may regulate structural plasticity in an activity-dependent manner. Taken together, we hypothesized that Rem2 could be a key regulator of cortical plasticity mechanisms in the intact nervous system.

Although Rem2 exhibits many hallmarks of a major activity-dependent plasticity molecule, it remains unclear if Rem2 actually regulates the functional responses of neural circuits in an experience-dependent manner, and, if so, which cellular and circuit mechanisms underlie these changes. Toward this
goal, we generated *Rem2* knockout mice to directly assess how Rem2 influences plasticity in the mammalian visual system. We found that *Rem2*^-/-^ mice exhibit a deficit in late-phase OD plasticity, whereas early-phase OD plasticity and adult OD plasticity are normal. These functional deficits depend specifically on deletion of *Rem2* from excitatory neurons in the cortex. At the cellular level, we found that *Rem2*^-/-^ neurons exhibit altered regulation of synaptic scaling and intrinsic excitability. Further, using sparse deletion methods, we found that the cell intrinsic effects on intrinsic excitability are cell-autonomous, and precede the effects on synaptic function. These data suggest that Rem2 provides a unique genetic inroad to understanding the fundamental processes that regulate intrinsic excitability at the molecular level, an area that has until now remained largely unexplored. Further, these data confirm that Rem2 operates at the nexus of plasticity pathways that regulate neuronal circuit morphology and connectivity in a manner that has important implications for circuit function.

## RESULTS

### Rem2 is expressed in cortex during the critical period

We first sought to determine when Rem2 is expressed in the developing rodent cortex based on the hypothesis that if Rem2 is an important molecule in activity-dependent plasticity pathways, then its expression should be modulated by neural activity evoked by natural sensory stimulation. Toward this end, rat cortical lysates were harvested at different developmental time points (from postnatal day (P) 1 to adult). Samples were analyzed by immunoblotting using an antibody that specifically recognizes Rem2 (Fig. 1A). We found that Rem2 expression peaks in the cortex around the time of eye opening (P9-P14, eyes open at P13 in rodents), and declines near the end of the critical period (P35, Gordon and Stryker, 1996). This result suggests that Rem2 is expressed during the developmental window in which robust synapse formation and activity-dependent refinement of cortical circuits occurs.

**Figure 1.**
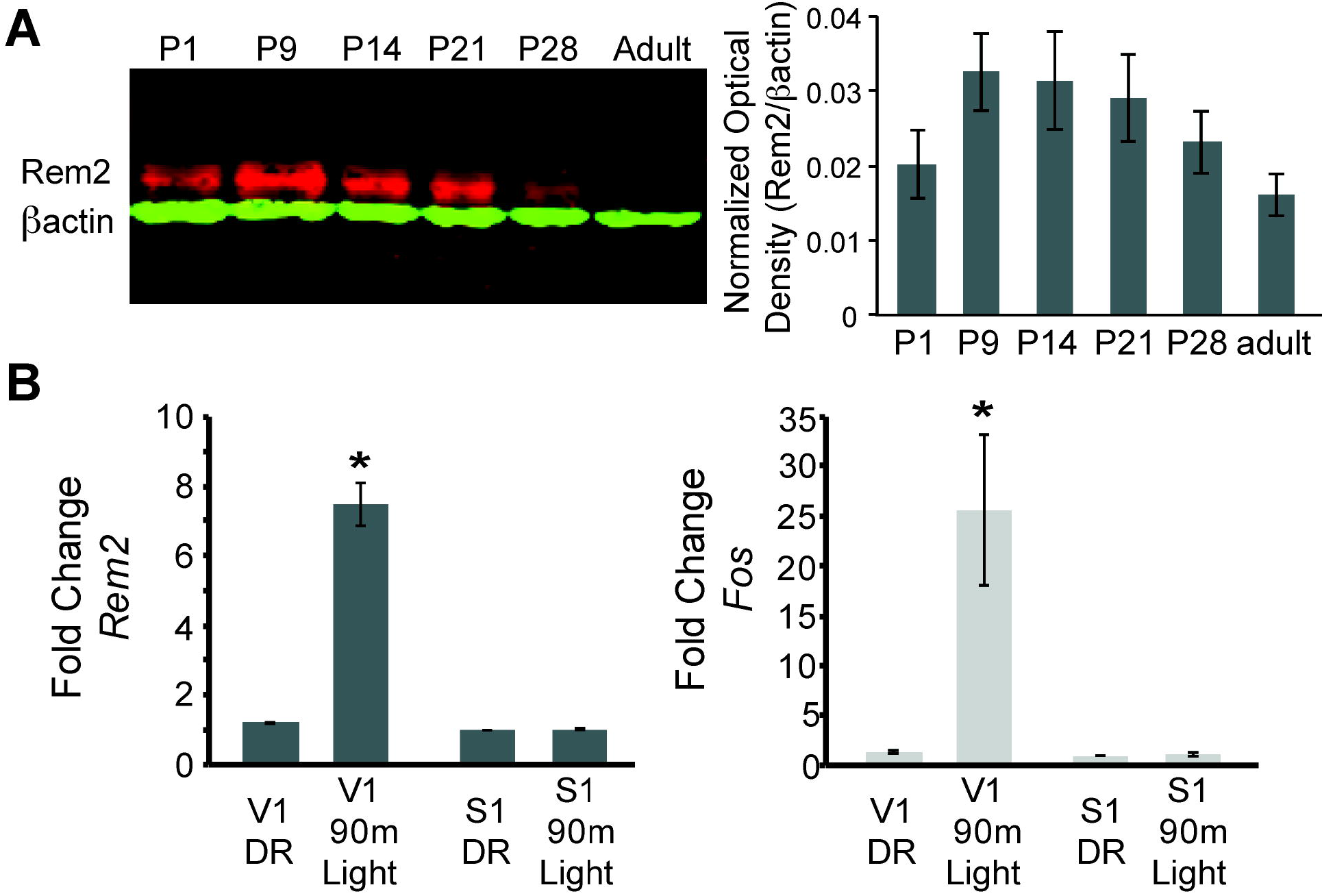
Rem2 expression is developmentally regulated and activity-dependent. **A**) (left) Western Blot of cortical lysate from rat brains at different developmental ages detected using anti-Rem2 (1:500); anti-Pactin (1:5000) was used as a loading control. (right) Quantification of relative Rem2 intensity at different developmental ages normalized to anti-βactin. **B**) Fold change in *Rem2* (left) or *Fos* mRNA expression (right) in isolated primary visual cortex (V1) or somatosensory cortex (S1) from P28 mice raised in the dark (DR, from P9-P28) or raised in the dark (P9-P28) and exposed to light for 90 m (90m Light). N=3 biological replicates of 4 mice in each experiment. mRNA levels were normalized to *Actb* levels and then to S1 DR condition and presented as mean ± SEM. * p < 0.05 from V1 DR by one-way ANOVA followed by a Dunnett’s post hoc test.

Given that Rem2 is expressed around eye opening (Fig. 1A) and that *Rem2* mRNA is modulated by neuronal activity (Ghiretti et al., 2014) we next asked if *Rem2* mRNA expression is regulated in mammalian visual cortex by sensory experience. To address this question, young mice were dark reared (DR) from P9 (prior to eye opening at P13) to P28 (during the peak of the critical period). At P28, one group of mice was exposed to 90 minutes of light stimulation while a separate group was kept in the dark. Following visual stimulation, primary visual cortex (V1) and primary somatosensory cortex (S1), used as a negative control, were micro-dissected and cDNA was prepared from total RNA for use in RT-qPCR experiments (Fig. 1B). Compared to DR littermates, 90 minutes of light stimulation led to a significant and rapid increase in *Rem2* mRNA expression in V1 with no change in *Rem2* mRNA expression in S1 (Fig 1B, left). As expected, we also observed an increase in mRNA levels of the immediate early gene *Fos* in response to changes in visual experience (Fig 1B, right). These data illustrate that Rem2 expression in V1 is modulated by visual stimulation coincident with experience-dependent development of visual cortical circuits. Interestingly, although our results demonstrate a reduction in baseline Rem2 protein expression around P28 and in adulthood (Fig. 1A, compared to P9 and P14), *Rem2* mRNA expression can be modulated by light stimulation both at P28 (Fig. 1B) and in the adult visual cortex (Mardinly et al., 2016). Thus, while baseline Rem2 levels may decline with age, Rem2 expression continues to be regulated by neuronal activity in discrete cell types into adulthood, suggesting that Rem2 function may be relevant throughout the life of the animal.

### Generation and validation of the *Rem2* null and conditional knockout mouse

To determine if Rem2 functions to regulate visual system plasticity *in vivo*, we generated a *Rem2* knockout mouse. Embryonic stem cell lines harboring a cassette with conditional potential (Fig. 2A, cassette strategy originally developed by Skarnes et al., 2011) at the *Rem2* locus were acquired from EUCOMM (Ref ID: 92501) and injected into blastocysts from which we obtained chimeras and germline transmission of this *Rem2* allele (referred to as *Rem2*^-/-^; Fig 2A middle). We verified correct insertion of the cassette at the *Rem2* locus both by Southern blot analysis (Fig. 2B) and by extensive PCR followed by sequencing (Fig. 2D, top). We also demonstrated that, as expected (Skarnes et al., 2011), insertion of the cassette disrupts Rem2 expression by immunoblotting (Fig. 2C), confirming a null allele at the protein level. The *Rem2*^-/-^ mice were also crossed with mice expressing Flp recombinase in the germline (JAX 009086), which produced a “floxed” (i.e. flanked by loxP sites) *Rem2* allele (referred to as *Rem2*^*flx*/*flx*^; Fig. 2A, bottom). The location of the loxP sites was confirmed by extensive PCR followed by sequencing across the entire targeted *Rem2* locus (Fig. 2D, bottom).

**Figure 2.**
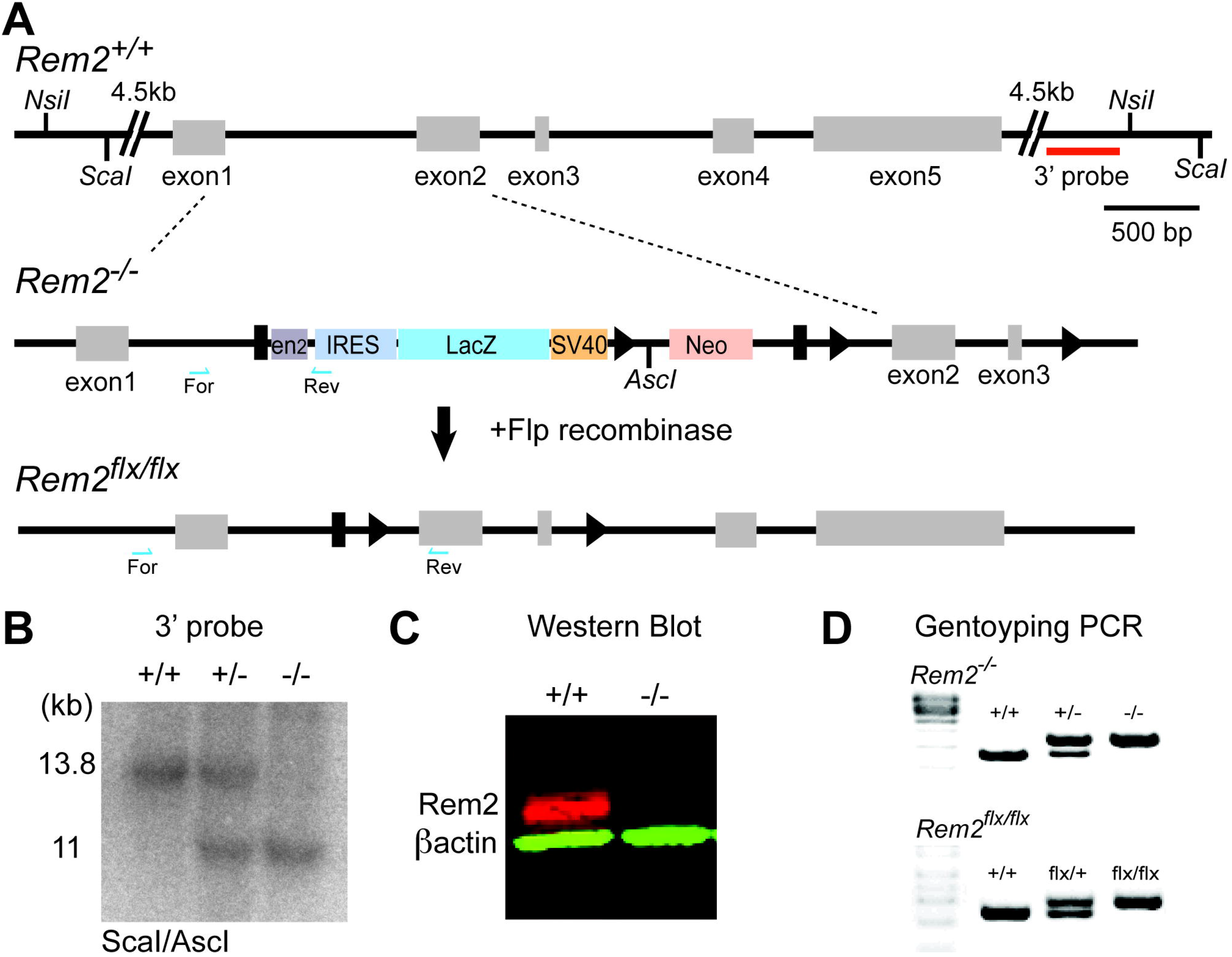
Generation of a Rem2 knockout mouse. **A**) The *Rem2* locus (top, *Rem2*^+/+^) with exons 1-5 depicted as gray boxes. Homologous recombination was used to insert a 7.5 kb cassette containing a mouse *En2* splice acceptor sequences (EN2), IRES, a LacZ gene, a SV40 polyadenylation, and a Neo gene flanked by FRT sites (black rectangles) and LoxP sites flanking exons 2 and 3 (indicated by black triangles). Insertion of this cassette yielded a *Rem2* null allele (Rem2^−/−^). Following germline transmission of the insert allele, the mice were crossed to a mouse expressing Flp recombinase in the germline, which produced a floxed *Rem2* allele (termed *Rem2*^*flx*/*flx*^, bottom). Blue half arrows indicate PCR forward and reverse primer locations. Scale bar, 500 bp. **B**) Southern blot analysis of ScaI/AscI-digested genomic DNA from mice that are *Rem2* wildtype (+/+), heterozygous (+/−) or homozygous (−/−) using the 3’ probe (red line, top A). Fragments of the predicted size (11 kb) indicate correct targeting. **C**) Western blot analysis of *Rem2*^+/+^ and *Rem2*^-/-^ mice confirming the loss of Rem2 expression. **D**) Genotyping PCR products of genomic DNA isolated from tails of mice that were (top) wildtype (*Rem2*^+/+^), heterozygous (*Rem2*^+/−^) or homozygous (*Rem2*^-/-^) or (bottom) wildtype (*Rem2*^+/+^), heterozygous (*Rem2*^*flx*/+^) or homozygous (Rem2^*flx*/*flx*^). See methods for genotyping details.

*Rem2*^-/-^ mice are viable and fertile and do not display any overt phenotypes. The genotypes of pups obtained from a *Rem2*^+/−^ cross are recovered in the expected Mendelian ratio. We closely examined the brains of these animals at different developmental ages (P7, P16, P21, and P30) to determine if loss of Rem2 resulted in any overt changes in brain or cortical structure. We found no difference in the brain/body weight ratio between *Rem2*^-/-^ and *Rem2*^+/+^ (i.e. wildtype) littermates at P7, P21 or P30 (Fig. S1A). In addition, we investigated the cortical thickness of the visual cortex at the aforementioned developmental ages. Visual cortex was identified using anatomical landmarks and measured from the deepest extent of layer 6 to the pial surface (just above the corpus callosum, Fig. S1D, blue line). While we found no significant change in cortical thickness at P7 and P21, we did observe a small increase in cortical thickness at P30 (Fig. S1D, p=0.05). We also examined cortical layer thickness by Nissl staining 30 μm sections through the visual cortex at P16 and found that layer thickness was indistinguishable between *Rem2*^-/-^ and wildtype mice (Fig. S1B, C). Thus, we conclude that cortical development at the anatomical level proceeds relatively normally despite the absence of Rem2.

A number of studies demonstrated that overexpression of Rem2 and other RGK proteins inhibit voltage-gated calcium channel function in a variety of cell types (Chen et al., 2005; Correll et al., 2008; Moore et al., 2013). Therefore, we sought to determine if resting calcium levels were perturbed by loss of *Rem2*. We prepared neuronal cultures from dissociated cortices obtained from E18 wildtype, *Rem2*^+/−^, and *Rem2*^-/-^ mice. At 5 days *in vitro* (DIV5), neurons were loaded with the ratiometric Ca^2+^ indicator Fura-2 and images of unstimulated cells were obtained using an epifluorescence microscope (Fig. S1E). Interestingly, we found no difference in the resting Ca^2+^ levels of cortical neurons obtained from wildtype, *Rem2*^+/−^, or *Rem2*^-/-^ mice (Fig. S1E). These data suggest that either endogenous Rem2 does not alter the Ca^2+^ permeability at rest in young cortical neurons, or that in the absence of Rem2 expression, compensatory mechanisms exist to maintain normal resting calcium levels.

To further understand the implications of *Rem2* deletion in visual system function, we examined visual response tuning properties in wildtype and *Rem2*^-/-^ mice using two-photon calcium imaging (Fig. S1F-I). The calcium indicator Oregon Green BAPTA-AM was bulk-loaded into cells of wildtype or *Rem2*^-/-^ mice (P31/32) 150 μm below the surface of the exposed binocular region of primary visual cortex (V1b; Fig. S1F). Visual stimuli consisting of a series of gratings moving in different directions (0° to 360° at 45° steps) were delivered at random to each eye (Fig. S1G, black arrows). No differences in measures of orientation tuning (circular variance, Ringach et al., 2002), or in measures of direction selectivity (circular variance in direction space, Mazurek et al., 2014) were observed between wildtype and *Rem2*^-/-^ neurons (Fig. S1H, I), indicating that basic visual response tuning properties are normal in *Rem2*^-/-^ mice. Taken together, our data suggests that, despite the expression of Rem2 during this developmental window, deletion of *Rem2* does not interfere substantially with initial brain or circuit development in the embryonic or early postnatal period of mouse development.

### Rem2 is required for normal critical period OD plasticity

Given that Rem2 expression is acutely regulated by visual experience (Fig. 1B), we sought to determine if Rem2 plays a role in activity-dependent processes in the visual cortex following sensory deprivation by investigating ocular dominance (OD) plasticity. We assayed critical period OD plasticity in *Rem2*^-/-^ mice or wildtype littermate controls that underwent normal visual experience (typically reared, TR: 12h light/12 h dark cycle) or that experienced monocular deprivation (MD) by lid suture for 2 days to measure early-phase OD plasticity or in a separate group of animals MD for 6 days to measure late-phase OD plasticity (Fig. 3A top). Using optical imaging of intrinsic signals (ISI), we measured cortical responses in the binocular portion of the visual cortex (V1b) to stimuli presented to either the left or right eye (Fig. 3B, representative images). We computed the ocular dominance index (ODI) of each animal by taking the response to stimulation of the contralateral (C) eye minus the response to stimulation of the ipsilateral (I) eye, normalized to the sum of these responses [ODI = (C −I)/(C + I)].

**Figure 3.**
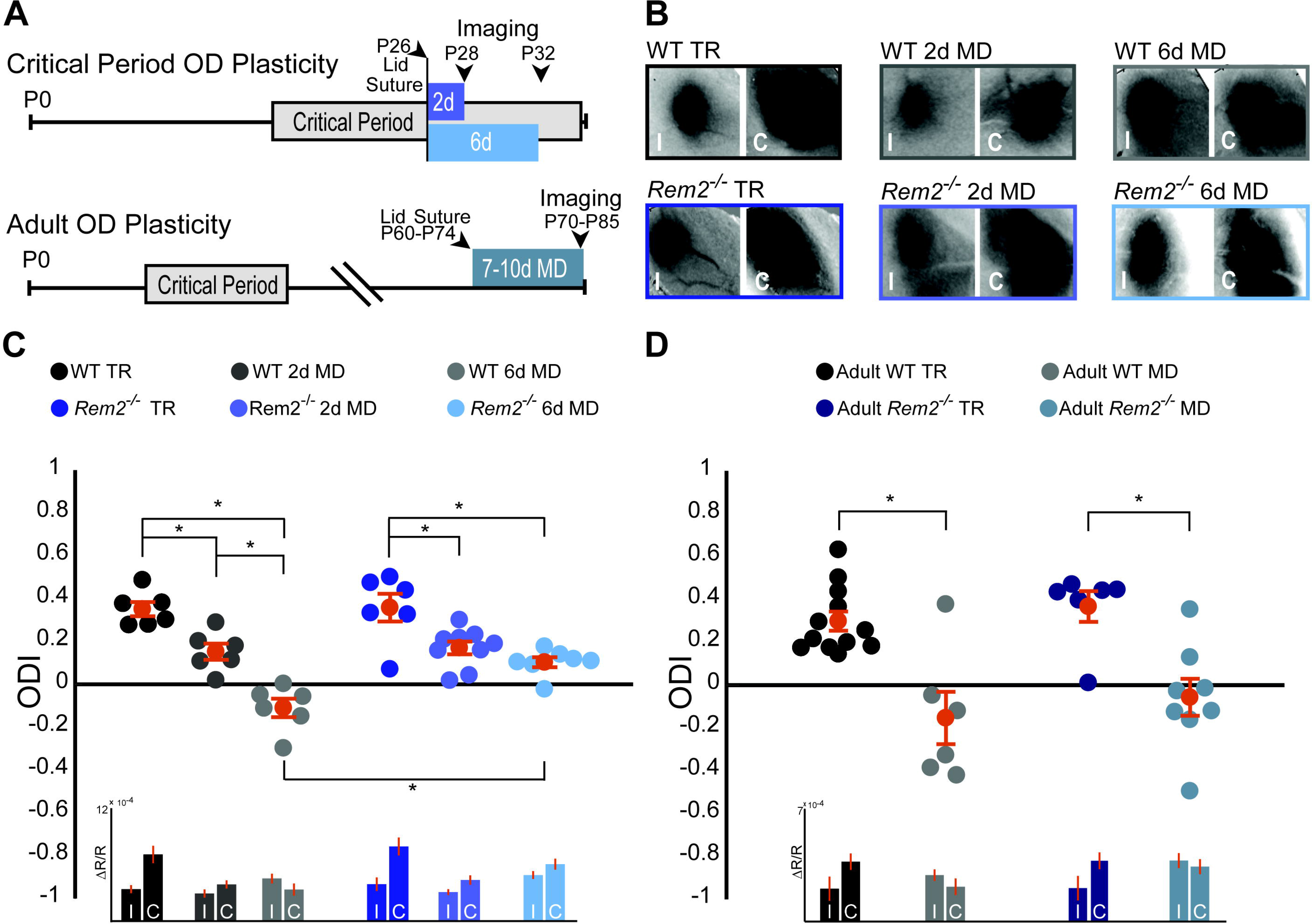
Rem2 is required for late-phase critical period ocular dominance plasticity. **A**) Representative experimental timeline for (top) critical period ocular dominance (OD) plasticity and (bottom) adult OD plasticity. **B**) Example fields showing intrinsic signal imaging data from the ipsilateral (I) and contralateral (C) eyes in wildtype or *Rem2*^-/-^ mice that were (left) typically reared (TR) or monocularly deprived (MD) for 2 days (middle) and 6 days (right). **C**) Ocular dominance index (ODI) for WT TR (black, n=6), WT 2d MD (dark gray, n=6), WT 6d MD (light gray, n=6), *Rem2*^-/-^ TR (blue, n=6), and *Rem2*^-/-^ 2d MD (medium blue, n=9), *Rem2*^-/-^ 6d MD (light blue, n=7). Mice are shown as circles for each animal. Orange circles with error bars represent the group averages. Inset at bottom: Changes in reflectance over baseline reflectance (ΔR/R) as measured by ISI driven ipsilateral (I) or contralateral (C) eye for wildtype and *Rem2*^-/-^ mice that were either typically reared or monocularly deprived. **D**) ODI for Adult WT TR (black, n=12), Adult WT MD (gray, n=6), Adult *Rem2*^-/-^ TR (navy blue, n=6), and Adult *Rem2*^-/-^ MD (gray blue, n=8). Inset: ΔR/R for Adult WT and *Rem2*^-/-^ TR and MD mice. Data is presented as mean ± SEM. *p < 0.05 by two-way ANOVA and Tukey post-hoc. Significance comparisons for ΔR/R in C and D insets are listed in Supplemental Table 1.

Wildtype typically reared littermate controls exhibited normal OD plasticity (Fig. 3C). Closure of the contralateral eye for 2 days (P26-28) produced an initial, significant shift in the ODI (Fig. 3C, WT TR = 0.35 ± 0.03 ODI; WT 2d MD = 0.15 ± 0.04 ODI, p < 0.01). In a separate experimental group, prolonged MD for 6 days (P26-32) resulted in a further robust shift in OD toward the ipsilateral eye (Fig 3C, WT 6d MD = −0.11 ± 0.04 ODI, p < 0.01 compared to WT TR) as has been previously reported (Gordon and Stryker, 1996; Heimel et al., 2007; Mrsic-Flogel et al., 2007). Consistent with prior reports (Frenkel and Bear, 2004; Gordon and Stryker, 1996; Heimel et al., 2007; Kaneko et al., 2008b; Mrsic-Flogel et al., 2007), responses to the contralateral (deprived) eye were decreased during the early-phase of deprivation (2d MD), while responses to the ipsilateral eye increased slightly during the late-phase of deprivation (6d MD; Fig. 3C, inset, Table S1).

*Rem2*^-/-^ mice with normal visual experience (*Rem2*^-/-^ TR) displayed an ODI similar to that observed in wildtype mice (Fig. 3C, *Rem2*^-/-^ TR = 0.35 ± 0.06 ODI, p=0.54 compared to WT TR). Closure of the contralateral eye in *Rem2*^-/-^ mice for 2 days with MD induced an initial, significant shift in the ODI similar to that observed in WT mice (Fig. 3C, *Rem2*^-/-^ 2d MD = 0.17 ± 0.03 ODI, p < 0.01), indicating that early-phase OD plasticity is intact in the absence of Rem2. As expected, the magnitude of depression of the deprived, contralateral eye responses following 2d MD in *Rem2*^-/-^ mice is similar to those observed in wildtype mice (Fig. 3C, inset 2d MD, Table S1), and suggests that a weakening of synaptic strength, reminiscent of LTD, is indeed induced in either the presence or absence of Rem2. However, in contrast to early phase plasticity, late-phase OD plasticity was altered in *Rem2*^-/-^ mice relative to their wildtype littermate controls (Fig. 3C). Following 6 days of MD, *Rem2*^-/-^ mice failed to display a further shift in ODI (Fig. 3C, *Rem2*^-/-^ 6d MD = 0.10 ± 0.02 ODI vs. WT 6d MD −0.11 ± 0.04 ODI, p<0.001 vs. *Rem2*^-/-^ 2d MD = 0.17 ± 0.03, p = 0.44), indicating a deficit in late-phase OD plasticity.

A closer examination of individual eye responses reveals interesting differences between WT and *Rem2*^-/-^ plasticity. Both WT and *Rem2*^-/-^ mice exhibited a decrease in responses to the contralateral (deprived) eye after 2d MD (Fig. 3C, inset). However, upon 6d of MD, WT mice exhibited an increase in ipsilateral (open) eye response (Fig 3C, inset WT 6d MD). This increased response has been previously attributed to homeostatic mechanisms (Kaneko et al., 2008; Lambo and Turrigiano, 2013), which presumably interact with continued reductions in responses to the contralateral eye to promote a further shift in ODI. By contrast, *Rem2*^-/-^ mice exhibited relatively equal increases in both the contralateral eye and ipsilateral eye responses with 6d MD. Thus, *Rem2*^-/-^ animals exhibited a non-competitive increase in activity during late-phase MD. These results raise the possibility that Rem2 may play an important role in regulating the absolute responsiveness of the cortex to visual stimulation, such as through regulation of neuronal excitability or synaptic scaling.

In order to gain more insight into the circuit mechanisms that might underlie the observed OD plasticity deficits of *Rem2*^-/-^ mice, we went on to examine adult OD plasticity. Unlike critical period OD plasticity, adult OD plasticity relies primarily on elimination of inhibitory synapses and does not require homeostatic synaptic scaling (Hofer et al., 2006; Lehmann and Lowel, 2008; Ranson et al., 2012; Sawtell et al., 2003; van Versendaal et al., 2012). To measure changes in adult OD plasticity, wildtype littermate controls or *Rem2*^-/-^ mice underwent normal visual experience (typically reared, TR: 12h light/12 h dark cycle) or monocular deprivation (MD) for 7-10 days between 10-12 weeks of age, a time well-beyond the classically defined critical period (Fig. 3A bottom), followed by ISI to measure cortical responses in V1b. Wildtype mice monocularly deprived for 7-10 days in adulthood produced a robust shift in ODI from a contralateral bias to an ipsilateral bias (Fig. 3D, Adult WT TR = 0.30 ± 0.04 ODI; Adult WT MD = −0.15 ± 0.12 ODI, p < 0.001). Similarly, *Rem2*^-/-^ mice that underwent monocular deprivation in adulthood also produced a significant shift in ODI following 7-10 days of MD (Fig. 3D, Adult *Rem2*^-/-^ TR = 0.37 ± 0.07 ODI; Adult *Rem2*^-/-^ MD = −0.05 ± 0.09 ODI, p < 0.01). Thus, taken together these data suggest that Rem2 is required specifically for late-phase critical period plasticity.

### Rem2 is required in cortical excitatory neurons for critical period OD plasticity

Ocular dominance plasticity during the critical period is dependent on the proper balance of network excitation and inhibition. To probe whether Rem2 regulates the plasticity of excitatory neurons, inhibitory neurons, or both, *Rem2*^*flx*/*flx*^ animals were crossed to mice directing Cre recombinase expression under the control of cell-type specific promoter elements. To assay the contribution of Rem2 expression in cortical excitatory pyramidal neurons we used the EMX1-Cre line (*EMX1*^*Cre*^, JAX #005628), where Cre expression is turned on early in embryonic development and is largely restricted to the dorsal telencephalon (Gorski et al., 2002). *Rem2*^+/+^; *EMX1*^*Cre*^ or *Rem2*^*flx*/*flx*^; *EMX1*^*Cre*^ mice were typically reared or monocularly deprived for 5-7 days as outlined above and ODI was measured using ISI. The *Rem2*^+/+^; *EMX1*^*Cre*^ mice showed a pronounced shift in their ODI following 5-7 days of MD as expected (Fig. 4A, *Rem2*^+/+^; *EMX1*^*Cre*^ TR = 0.33 ± 0.06 ODI; *Rem2*^+/+^; *EMX1*^*Cre*^ MD = −0.06 ± 0.06 ODI, p < 0.001). However, *Rem2* deletion specifically from excitatory, cortical neurons (*Rem2*^*flx*/*flx*^; *EMX1*^*Cre*^) resulted in diminished OD plasticity following MD (Figs. 4A, *Rem2*^*flx*/*flx*^; *EMX1*^*Cre*^ TR = 0. 31 ± 0.04 ODI; *Rem2*^*flx*/*flx*^; *EMX1*^*Cre*^ MD = 0.21 ± 0.06 ODI, p=0.61). We make two important conclusions based on this data. First, Rem2 is required in the cortex to mediate proper OD plasticity, as the EMX1 promoter does not drive Cre expression in other upstream regions of the visual system such as thalamus and retina (Gorski et al., 2002). Second, within the cortex, Rem2 is required in excitatory pyramidal neurons for critical period ocular dominance plasticity to occur.

**Figure 4.**
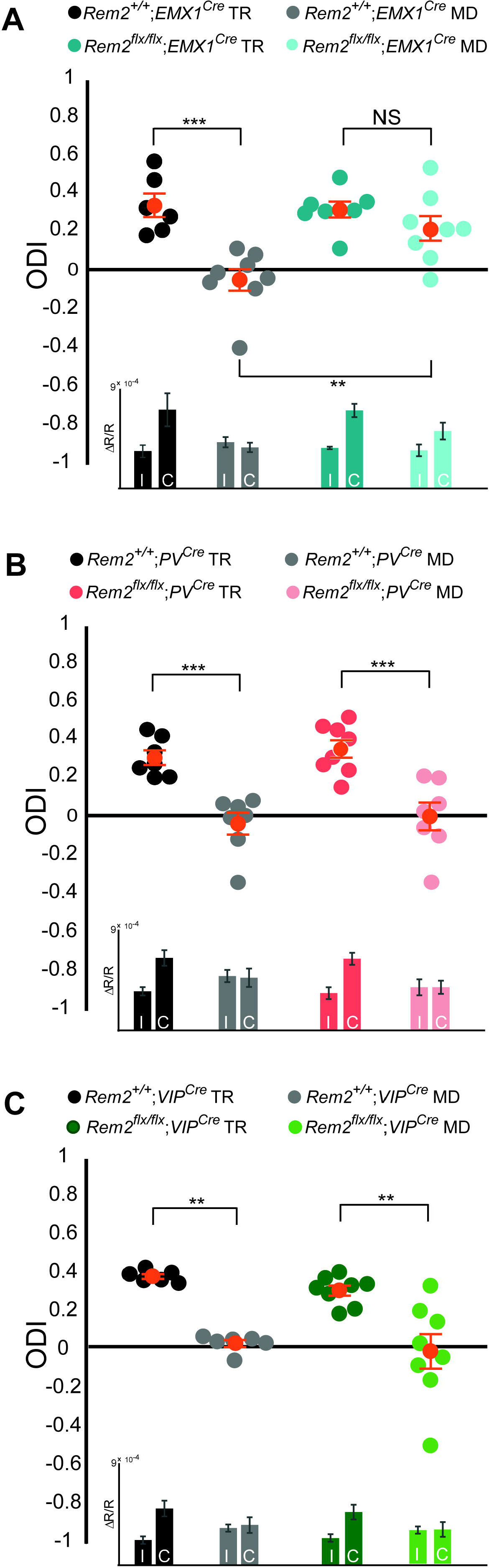
Rem2 is required in cortical excitatory neurons for ocular dominance plasticity. **A**) Ocular dominance index (ODI) for *Rem2*^+/+^; *EMX1*^*Cre*^ typically reared (TR, black, n=6), *Rem2*^+/+^; *EMX1*^*Cre*^ monocularly deprived (MD, gray, n=8), *Rem2*^-/-^; *EMX1*^*Cre*^ TR (dark teal, n=7) or *Rem2*^-/-^; *EMX1*^*Cre*^ MD (light teal, n=8). Inset: ΔR/R for *Rem2*^+/+^; *EMX1*^*Cre*^ and *Rem2*^-/-^; *EMX1*^*Cre*^ TR and MD mice. **B**) ODI for *Rem2*^+/+^; *PV*^*Cre*^ TR (black, n=7), *Rem2*^+/+^; *PV*^*Cre*^ MD (gray, n=7), *Rem2*^-/-^; *PV*^*Cre*^ TR (magenta, n=8) or *Rem2*^-/-^; *PV*^*Cre*^ MD (light pink, n=7). Inset: ΔR/R for *Rem2*^+,+^; *PV*^*Cre*^ and *Rem2*^-/-^; *PV*^*Cre*^ TR and MD mice. **C**) ODI for *Rem2*^+/+^; *VIP*^*Cre*^ TR (black, n=6), *Rem2*^+/+^; *VIP*^*Cre*^ MD (gray, n=6), *Rem2*^-/-^; *VIP*^*Cre*^ TR (green, n=8) or *Rem2*^-/-^; *PV*^*Cre*^ MD (light green, n=8). Inset: ΔR/R for *Rem2*^+,+^; *VIP*^*Cre*^ and *Rem2*^-/-^; *VIP*^*Cre*^ TR and MD mice. Each animal is depicted as an individual circle. Orange circles with error bars represent the averages for each group ± SEM. Data is presented as mean ± SEM. *p < 0.05 by two-way ANOVA with Tukey post-hoc. Significance comparisons for ΔR/R in C and D insets are listed in Supplemental Table 1.

It is well established that inhibitory tone plays a critical role in opening and closing the critical period for OD plasticity (Hensch and Fagiolini, 2005; Kuhlman et al., 2013; Southwell et al., 2010; van Versendaal and Levelt, 2016). Therefore, we sought to determine whether Rem2 expression in inhibitory interneurons contributes to OD plasticity. To this end, we crossed the *Rem2*^*flx*/*flx*^ mice to a mouse line expressing Cre recombinase under control of the parvalbumin (PV) promoter (*PV*^*Cre*^, JAX#017320) as PV+ interneurons are one of the most abundant interneuron subtypes in the cerebral cortex and disinhibition from interneurons plays an important role in OD plasticity (Butt et al., 2005; Kuhlman et al., 2013; Rudy et al., 2011). As expected, we observed a significant shift in the ODI in *Rem2*^+/+^; *PV*^*Cre*^ mice following monocular deprivation (Figs. 4B, *Rem2*^+/+^; *PV*^*Cre*^ TR = 0.30 ± 0.04 ODI; *Rem2*^+/+^; *PV*^*Cre*^ MD = −0.05 ± 0.06 ODI, p=0.001). However, in contrast to the lack of OD plasticity observed in the *Rem2*^*flx*/*flx*^; *EMX1*^*Cre*^ mice, deletion of *Rem2* from PV+ interneurons (*Rem2*^*flx*/*flx*^; *PV*^*Cre*^) had no effect on OD plasticity (Figs. 4B, *Rem2*^fIx/fIx^; *PV*^*Cre*^ TR = 0.34 ± 0.05 ODI, *Rem2*^*flx*/*flx*^; *PV*^*Cre*^ MD = −0.01 ± 0.07 ODI, p<0.001). These data demonstrate that Rem2 is not required in PV^+^ interneurons for OD plasticity.

Vasoactive intestinal peptide (VIP) positive interneurons regulate cortical gain control during arousal (Fu et al., 2014), are integral to a disinhibitory circuit that enhances adult OD plasticity (Fu et al., 2015), and regulate visual acuity in an experience-dependent manner (Mardinly et al., 2016). Additionally, previous work demonstrated a significant increase in *Rem2* mRNA expression in VIP^+^ interneurons following light exposure in dark reared mice (Mardinly et al., 2016). To assay whether Rem2 is required in VIP^+^ interneurons for critical period plasticity we crossed the *Rem2*^*flx*/*flx*^ mice to a mouse line expressing Cre recombinase under control of the VIP promoter (*VIP*^*Cre*^, JAX#010908). We observed a significant shift in the ODI in *Rem2*^+/+^; *VIP*^*Cre*^ mice following monocular deprivation (Figs. 4C, *Rem2*^+/+^; *VIP*^*Cre*^ TR = 0.39 ± 0.01 ODI; *Rem2*^+/+^; *VIP*^*Cre*^ MD = 0.04 ± 0.02 ODI, p=0.01). A similar shift was observed in the monocularly deprived *Rem2*^*flx*/*flx*^; *VIP*^*Cre*^ mice, indicating that deletion of *Rem2* from VIP^+^ interneurons has no effect on OD plasticity (Figs. 4C, *Rem2*^*flx*/*flx*^; *VIP*^*Cre*^ TR = 0.32 ± 0.03 ODI; *Rem2*^*flx*/*flx*^; *VIP*^*Cre*^ MD = −0.00 ± 0.09 ODI, p<0.01). Taken together these data demonstrate that Rem2 is not required in either PV^+^ or VIP^+^ interneurons for critical period OD plasticity.

To further examine inhibition in the cortex of the *Rem2*^-/-^ animals, we assayed miniature inhibitory postsynaptic current (mIPSC) amplitude and frequency in layer 2/3 pyramidal neurons (Fig. S2). We found no change in mIPSC frequency (Fig. S2A, B; WT TR = 5.63 ± 0.09 Hz; WT 6d MD = 5.31 ± 0.13 Hz, p=0.931) or amplitude (Fig. S2C, D; WT TR = −31.04 ± 1.0 pA; WT 6d MD = −31.92 ± 0.92 pA, p=0.852) in response to 6 days of MD in wildtype neurons. Additionally, mIPSC frequency and amplitude recorded from neurons in *Rem2*^-/-^ mice that were typically reared was similar to WT TR neurons, suggesting that deletion of *Rem2* does not alter baseline inhibitory synapse formation or transmission (Fig. S2, (B) *Rem2*^-/-^ TR frequency = 5.27 ± 0.13 Hz; (C, D) amplitude = −29.22 ± 1.47 pA, p=0.73 compared to WT TR). We did, however, observe a significant increase in mIPSC frequency, but not amplitude, in *Rem2*^-/-^ mice with 6 days of MD (Fig. S2B-D, *Rem2*^-/-^ 6d MD frequency = 6.32 ± 0.13 Hz, p=0.03; *Rem2*^-/-^ 6d MD amplitude = −29.02 ± 1.0 pA, p=0.35 compared to *Rem2*^-/-^ TR), suggesting an increase in inhibitory tone perhaps in response to increased cortical excitability. This result, combined with our ODI data in the *Rem2*^*flx*/*flx*^; *PV*^*Cre*^ mice and *Rem2*^*flx*/*flx*^; *VIP*^*Cre*^, suggests that Rem2 plays a minor role in regulating cortical inhibition specifically in response to prolonged MD, perhaps in response to altered excitability.

### Rem2 is necessary for homeostatic synaptic scaling

At the circuit level, monocular deprivation causes biphasic changes in cortical excitability, which underlies the observed shift in ocular dominance. Early-phase OD plasticity (1-3d MD) results in an initial decrease in responsiveness to the closed eye largely dependent on LTD-like mechanisms (Cooke and Bear, 2014; Crozier et al., 2007; Yoon et al., 2009), while the slower gain of responsiveness to the open eye during late-phase OD plasticity (4-6d MD) is due to homeostatic mechanisms including postsynaptic scaling up of excitatory synapses and intrinsic excitability homeostasis (Kaneko et al., 2008b; Lambo and Turrigiano, 2013; Mrsic-Flogel et al., 2007; Smith et al., 2009). The presence of a late-phase ocular dominance plasticity phenotype, which resulted in enhanced visual responsiveness at 6d MD in *Rem2*^-/-^ animals, led us to question whether homeostatic mechanisms might be enhanced or altered in the absence of Rem2. To address this possibility, we examined the synaptic and intrinsic properties of layer 2/3 pyramidal neurons using *ex vivo* slice experiments following either 2 days or 6 days of monocular deprivation.

To begin, we assayed changes in postsynaptic inputs by measuring miniature excitatory postsynaptic current (mEPSC) amplitudes. Both wildtype and *Rem2*^-/-^ littermates were either typically reared or monocularly deprived for 2 or 6 days starting at P26 and continuing until P28 or P32, respectively (Fig. 5A, B; representative traces). Whole-cell voltage-clamp recordings were used to measure mEPSCs in layer 2/3 pyramidal neurons in acute slices. We found that 2 days of MD in wildtype layer 2/3 pyramidal neurons caused a significant decrease in mEPSC amplitude (Fig. 5A, C and E (left); WT TR = −9.8 ± 0.1 pA; WT 2d MD = −8.87 ± 0.08 pA, p=0.015) as previously reported (Lambo and Turrigiano, 2013). Similarly, recordings from cortical slices obtained from *Rem2*^-/-^ mice with 2 days of MD also resulted in a significant decrease in mEPSC amplitude (Fig. 5A, C and E (left), *Rem2*^-/-^ P28 TR = −9.76 ± 0.08 pA; *Rem2*^-/-^ 2d MD = −9.12 ± 0.08 pA, p=0.014). This data, in agreement with our 2d MD ODI data in *Rem2*^-/-^ mice (Fig. 3C), suggests that in the absence of Rem2, layer 2/3 cortical neurons are sensitive to the decrease in drive that occurs as a result of MD and decrease their excitatory postsynaptic strength as expected (Lambo and Turrigiano, 2013).

**Figure 5.**
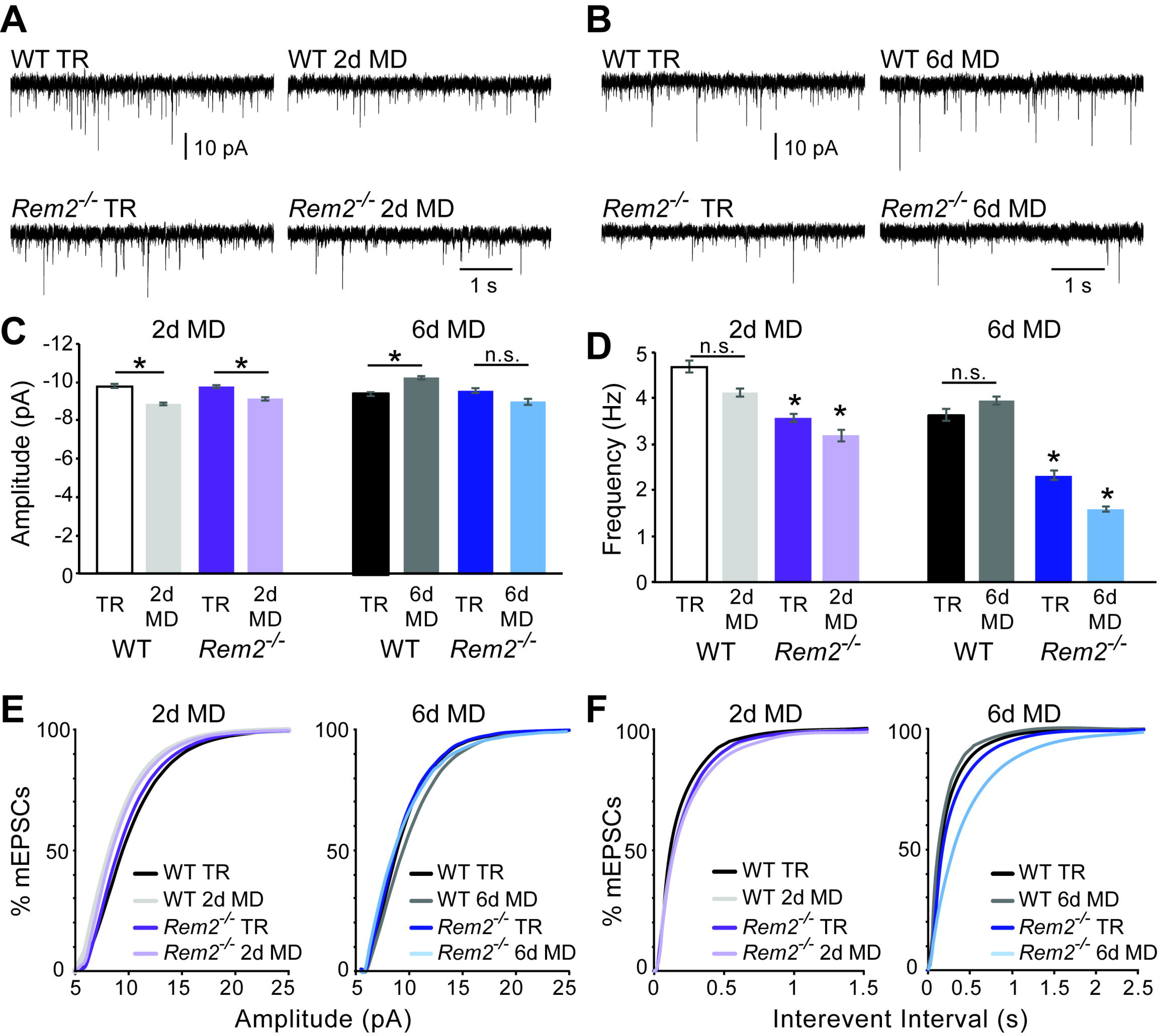
Rem2 is required for postsynaptic strengthening following 6 days of monocular deprivation. **A**) Representative whole-cell voltage clamp recordings of mEPSCs from layer 2/3 pyramidal neurons in binocular visual cortex (V1b) of wildtype typically reared mice at P28 (WT TR), wildtype mice undergoing 2 days of monocular deprivation (WT 2d MD), *Rem2*^-/-^ typically reared mice at P28 (*Rem2*^-/-^ TR), or *Rem2*^-/-^ mice following 2 days of monocular deprivation (*Rem2*^-/-^ 2d MD). **B**) Representative voltage-clamp traces from layer 2/3 pyramidal neurons from V1b of wildtype typically reared mice at P32 (WT TR), wildtype mice undergoing 6 days of monocular deprivation from P26-P32 (WT 6d MD), *Rem2*^-/-^ typically reared mice at P32 (*Rem2*^-/-^ TR) or following 6 days of MD (*Rem2*^-/-^ 6d MD). **C**) Average mEPSC amplitude recorded from wildtype or *Rem2*^-/-^ layer 2/3 pyramidal neurons with normal visual experience at (left) P28 (WT TR, white, n=25; *Rem2*^-/-^ TR, dark purple, n=29), or following 2 days of monocular deprivation (WT DR, light gray, n=25; *Rem2*^-/-^ 2d MD, light purple, n=23) or (right) at P32 with normal visual experience (WT TR, black, n=24; *Rem2*^-/-^ TR, dark blue, n=24) or following 6 days of monocular deprivation (WT 6d MD, gray, n=25; *Rem2*^-/-^ 6d MD, light blue, n=25). n=4 animals per experimental condition. **D**) Average mEPSC frequency in wildtype and *Rem2*^-/-^ mice undergoing 2 days (Left) or 6 days (Right) of monocular deprivation compared to typically reared age-matched controls. **E**) Cumulative distribution plot of mEPSC amplitude or (**F**) Interevent Interval recorded in wildtype and *Rem2*^-/-^ mice following 2 days (Left) or 6 days (Right) of monocular deprivation. Data is presented as mean ± SEM. *p < 0.05, by two-way ANOVA and Tukey post-hoc for mEPSC frequency and amplitude mean data plots.

We next examined the consequence of 6 days of MD in the absence of Rem2 by again assaying mEPSC amplitude. We hypothesized that if *Rem2*^-/-^ neurons were able to undergo homeostatic postsynaptic scaling, we would observe an increase in mEPSC amplitude, as previously reported in rat cortex (Lambo and Turrigiano, 2013). While we observed a significant increase in mEPSC amplitude in neurons in slices obtained from wildtype animals following 6 days of MD (Fig. 5B, C and E (right); WT TR = −9.46 ± 0.1 pA; WT 6d MD = −10.31 ± 0.09 pA, p=0.024), we failed to observe a significant increase in mEPSC amplitude in neurons from *Rem2*^-/-^ visual cortex following 6 days of MD (Fig. 6B, C and E (right); *Rem2*^-/-^ TR = −9.64 ± 0.12 pA; *Rem2*^-/-^ 6d MD = −9.04 ± 0.14 pA, p=0.535). Thus, these data demonstrate that excitatory postsynaptic scaling up is aberrant in *Rem2*^-/-^ mice.

**Figure 6.**
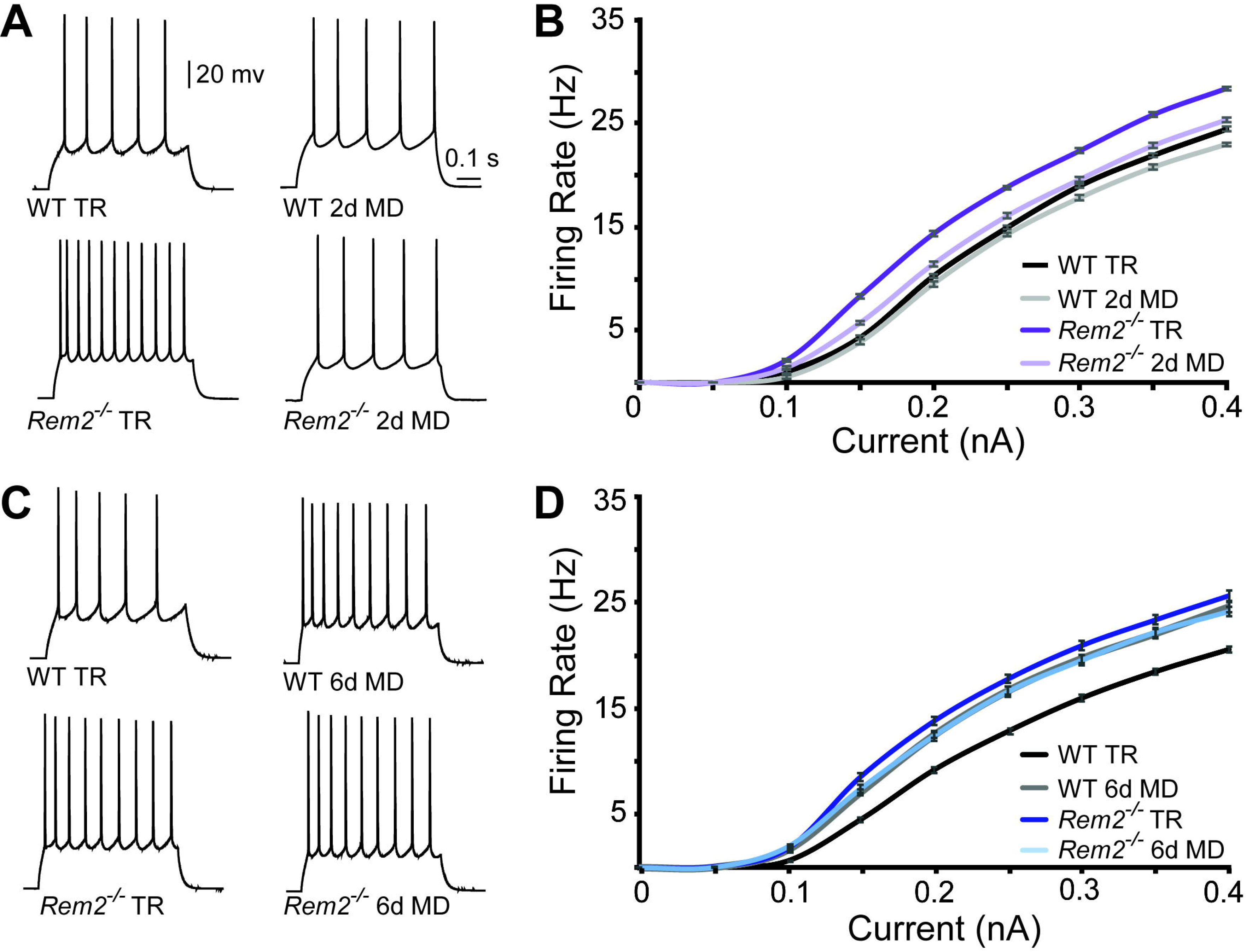
Rem2 alters the intrinsic excitability of layer 2/3 pyramidal neurons. **A**) Example traces of evoked action potential responses (0.2 nA current injected) of neurons from WT or *Rem2*^-/-^ mice either typically reared (TR) to P28 or monocularly deprived for 2 days from P26-P28 (2d MD). **B**) Average *f-I* curves for WT TR (black line, n=25), WT 2d MD (gray line, n=22), *Rem2*^-/-^ TR (dark purple line, n=23) and *Rem2*^-/-^ 2d MD (light purple line, n=22) in response to normal visual experience or 2 days of monocular deprivation. n=4 animals per experimental condition. **C**) Examples of evoked responses from neurons of WT or *Rem2*^-/-^ following at P32 that were either typically reared (WT TR and *Rem2*^-/-^ TR) or monocularly deprived for 6 days from P26-P32 (WT 6d MD and *Rem2*^-/-^ 6d MD). **D**) Average *f-I* curves for WT TR (black line, n=32), WT 6d MD (gray line, n=22), *Rem2*^-/-^ TR (dark blue line, n=24) and *Rem2*^-/-^ 6d MD (light blue line, n=22) in response to normal visual experience or after 6 days of monocular deprivation. n=5 mice for WT TR and 4 mice for all other conditions.

We also examined changes in mEPSC frequency in response to 2 or 6 days of MD. As expected, we found no change in mEPSC frequency in wildtype mice following either 2 days (Fig. 5D, F (left); WT TR = 4.69 ± 0.13 Hz; WT 2d MD = 4.12 ± 0.1 Hz, p=0.515) or 6 days (Fig. 5D, F (right); WT TR = 3.65 ± 0.13 Hz; WT 6d MD = 3.95 ± 0.1 Hz, p=0.551) of MD. Interestingly, both P28 and P32 typically reared *Rem2*^-/-^ mice displayed a significant decrease in mEPSC frequency when compared to their wildtype littermate controls reared under the same conditions (Fig. 5D, F; *Rem2*^-/-^ P28 TR = 4.13 ± 0.1 Hz, p=0.03; *Rem2*^-/-^ P32 TR = −2.34 ± 0.1 Hz, p=0.0005). The frequency of mEPSC events was still further decreased following both 2 days (Fig. 5E, F (left); *Rem2*^-/-^ 2d MD = 3.19 ± 0.12 Hz, p=0.003) and 6 days of MD (Fig. 5E, F (right); *Rem2*^-/-^ 6d MD = 1.6 ± 0.05 Hz, p<0.0001) in cortical neurons isolated from *Rem2*^-/-^ compared to wildtype animals. These changes in mEPSC frequency could reflect any number of differences between the cortical synaptic connections in the *Rem2*^-/-^ and wildtype mice including changes in excitatory synapse formation, presynaptic neurotransmitter release, or synapse maintenance and pruning mechanisms.

To determine if *Rem2* deletion affects excitatory synapse density in visual cortex, wildtype and *Rem2*^-/-^ mice were either light reared from birth (LR, i.e. housed in a 12hr light/12hr dark cycle) or dark reared (DR) from P9 (before eye opening) to P30 in a light-sealed dark box. At P30, the brains from both groups were harvested and Golgi histology performed. Spine density was calculated by counting the number of dendritic protrusions from a 50μm segment of dendrite per neuron imaged from a terminal branch tip off of the apical dendritic tree located 50 - 100 μm from the cell soma. In wildtype mice, dark rearing resulted in a significant reduction in spine density, similar to previous reports (Fig. S3: WT LR = 0.98 ± 0.01 spines/μm; WT DR = 0.83 ± 0.01 spines/μm, p<0.0001; (Valverde, 1971; Wallace and Bear, 2004). In the absence of Rem2 under normal visual experience (*Rem2*^-/-^ LR), spine density was significantly reduced compared to wildtype (Fig. S3; *Rem2*^-/-^ LR = 0.82 ± 0.02 spines/μm, p<0.0001 compared to WT LR). However, no further reduction of spine density was observed in *Rem2*^-/-^ mice following dark rearing (Fig. S3; *Rem2*^-/-^ DR = 0.87 ± 0.01 spines/μm, p=0.58 compared to *Rem2*^-/-^ LR). While these data do not rule out a presynaptic function for Rem2, they are consistent with the mEPSC data (Fig. 5) and suggest that one function of Rem2 is to promote experience-dependent synapse formation and function.

### Rem2 alters the homeostatic regulation of intrinsic excitability

While we did observe homeostatic deficits in synaptic scaling, these effects cannot explain the enhanced responsiveness observed in *Rem2*^-/-^ animals following 6d MD. Specifically, the lack of synaptic scaling up would suggest that the cortex should be less responsive to visual stimulation as opposed to more responsive as observed with ISI. We therefore turned our focus to intrinsic excitability, which is also homeostatically regulated in response to MD to counteract a perturbation in network drive (Desai et al., 1999; Lambo and Turrigiano, 2013; Maffei and Turrigiano, 2008; Marder and Goaillard, 2006; Turrigiano et al., 1998), and asked if layer 2/3 pyramidal neurons in *Rem2*^-/-^ mice were able to alter their intrinsic excitability following monocular deprivation.

To assess changes in intrinsic excitability, we measured the frequency of action potential firing in response to a series of depolarizing current steps (*f-I* curves) applied in the presence of pharmacological blockers of synaptic transmission (Fig. 6). Surprisingly, *Rem2*^-/-^ mice exhibited a pronounced leftward shift of the *f-I* curve under normal rearing conditions (Fig. 6), suggesting that at least one function of Rem2 is to maintain the intrinsic excitability of the neuron. This shift occurred in the absence of changes in R_IN_, V_m_ or C_m_ (Table S2) between wildtype and *Rem2*^-/-^ neurons, although we did observe a nonsignificant trend towards increased R_IN_ in *Rem2*^-/-^ neurons (at P32: WT 104.15 ± 1.64 GΩ; *Rem2*^-/-^ 112.06 ± 1.62 GΩ, see Table S2). Following 2 days of MD, wildtype neurons displayed no change in their *f-I* curve (Fig. 6A-B, compare WT TR to WT 2d MD), while *Rem2*^-/-^ neurons displayed decreased firing (Fig. 6A-B, compare *Rem2*^-/-^ TR to *Rem2*^-/-^ 2d MD), such that the *f-I* curves obtained from *Rem2*^-/-^ neurons were indistinguishable from wildtype (Fig. 6B, compare *Rem2*^-/-^ 2d MD to WT TR). Thus, while under normal wildtype conditions no change in intrinsic excitability is observed in response to 2 days of MD, in the absence of Rem2, neurons aberrantly respond to decreased network drive (e.g. weakened synaptic strength) by decreasing their intrinsic excitability.

When examined at 6 days of MD, wildtype neurons shifted their *f-I* curve to the left (Fig. 6C-D, compare WT TR to WT 6d MD), indicating a homeostatic increase in intrinsic excitability. However, neurons from *Rem2*^-/-^ mice failed to further increase their intrinsic excitability in response to 6 days of MD (Fig. 6D, compare *Rem2*^-/-^ TR to *Rem2*^-/-^ 6d MD). One interpretation of this data is that Rem2 is required for homeostatic increases, but not decreases in intrinsic excitability. Alternatively, it could be that in the absence of Rem2 homeostatic increases in intrinsic excitability are occluded by the already shifted “baseline-level” of excitability.

These data provide a parsimonious explanation for the intrinsic signal imaging results observed in *Rem2*^-/-^ animals (Fig. 3). At least part of the decrease in responsiveness observed with 2d MD in *Rem2*^-/-^ mice can be explained by the rightward shift in intrinsic excitability compared to *Rem2*^-/-^ TR mice (Fig. 6B). Additionally, the non-competitive increase in responsiveness observed following 6d MD could reflect the increased intrinsic excitability observed *in vitro* (Fig. 6D). These data support the conclusion that Rem2 regulates the absolute responsiveness of the cortex to visual stimulation through homeostatic adjustments in intrinsic excitability.

### Rem2 functions cell-autonomously to regulate intrinsic excitability

Given the unexpected increase in intrinsic excitability observed in TR *Rem2*^-/-^ mice, we sought to determine whether this shift depended on circuit level changes in network excitability as a result of synapse loss, or rather reflected a cell-autonomous function of Rem2. To address this question, we performed acute, sparse deletion of *Rem2* in layer 2/3 pyramidal neurons in binocular visual cortex. *Rem2*^*flx*/*flx*^ mice were injected at P18-20 with either a dilute control virus (AAV-GFP) or virus expressing Cre recombinase (AAV-Cre-GFP; Fig. 7A). Our injection strategy causes expression of GFP, or expression of GFP and deletion of *Rem2*, from a few dozen neurons, while leaving the majority of the circuit unaffected. We have previously verified using qPCR that 3-5 days of viral infection is sufficient to delete *Rem2* exons 2 and 3 (Kenny et al., 2017, Fig. 1A). Acute cortical slices were then isolated either 4 days post infection (4 d.p.i., Fig. 7A left) or 10-12 d.p.i. (Fig. 7A right), and *f-I* curves constructed from GFP^+^, layer 2/3 pyramidal neurons to determine how acute deletion of *Rem2* affects neuronal intrinsic excitability.

Surprisingly, we found that acute, sparse deletion of *Rem2* from layer 2/3 pyramidal neurons 4 d.p.i. led to a significant increase in intrinsic excitability in response to current injection (Fig. 7B, *Rem2*^*flx*/*flx*^ + AAV-GFP-Cre 4 d.p.i.) compared to those with normal Rem2 expression (Fig. 7B, *Rem2*^*flx*/*flx*^ + AAV-GFP 4 d.p.i.). Similarly, deletion of *Rem2* for 10-12 d.p.i. also resulted in a significant increase in intrinsic excitability (Fig. 7C, *Rem2*^*flx*/*flx*^ + AAV-GFP-Cre 10-12 d.p.i.) compared to their wildtype littermate controls (Fig. 7C, *Rem2*^*flx*/*flx*^ + AAV-GFP 10-12 d.p.i.). These results indicate that Rem2 can regulate intrinsic excitability in a cell-autonomous manner and in the absence of sensory manipulation.

However, this experiment does not resolve the confounding relationship between both altered intrinsic excitability and synapse function observed in the *Rem2*^-/-^ mice. Therefore, in order to determine whether the shift in intrinsic excitability was due to a preceding change in the number of functional synapses, we also measured mEPSC frequency and amplitude following 4 or 10-12 days of *Rem2* deletion. Interestingly, we found that acute deletion of *Rem2* measured 4 d.p.i. was not sufficient to significantly alter mEPSC frequency or amplitude (Fig. 7D, (left) frequency: *Rem2*^*flx*/*flx*^ + AAV-GFP = 5.59 ± 0.22 Hz; *Rem2*^*flx*/*flx*^ + AAV-GFP-Cre = 5.10 ± 0.24 Hz, p=0.423; (right) amplitude: *Rem2*^*flx*/*flx*^ + AAV-GFP = −10.28 ± 0.31 pA; *Rem2*^*flx*/*flx*^ + AAV-GFP-Cre = −9.60 ± 0.24 pA, p=0.386). Conversely, by 10-12 d.p.i. a significant decrease in mEPSC frequency (Fig. 7E, left; *Rem2*^*flx*/*flx*^ + AAV-GFP = 4.38 ± 0.15 Hz; *Rem2*^*flx*/*flx*^ + AAV-GFP-Cre = 2.92 ± 0.19 Hz, p<0.01) while no change in amplitude was observed (Fig. 7E right; *Rem2*^*flx*/*flx*^ + AAV-GFP = −9.76 ± 0.22 pA; *Rem2*^*flx*/*flx*^ + AAV-GFP-Cre = −10.42 ± 0.26 Hz, p=0.38).

Additionally, we quantified changes in spine density 10 d.p.i. by utilizing sparse injections of AAV-Cre-GFP into an Ai9 Cre reporter mouse line harboring a loxP-flanked STOP cassette preventing transcription of a CAG promoter-driven tdTomato (Ai9, JAX#007909) crossed to our conditional *Rem2*^*flx*/*flx*^ line. Spine density, head width, and neck length from both *Rem2*^+/+^; *TdT*^*flx*/*flx*^ + AAV-GFP-Cre and *Rem2*^*flx*/*flx*^; *TdT*^*flx*/*flx*^ +AAV-GFP-Cre mice was quantified using Reconstruct (Fiala, 2005). With acute deletion of *Rem2* (10 d.p.i.), we observed a trend toward decreased spine density and spine head width, as well as a statistically significant decrease in spine neck length (Fig. S4). These results, together with our mEPSC data (Fig. 7D-E), suggest that, approximately 10 days following *Rem2* deletion, there is a significant decrease in functional synapse number, prior to spine shrinkage and eventual removal. Therefore, we conclude that Rem2 functions in a cell-autonomous manner to regulate intrinsic excitability irrespective of excitatory synapse density. Thus, Rem2 is a novel regulator of intrinsic excitability in excitatory cortical neurons required to sculpt circuit function.

## DISCUSSION

The present study identifies the activity-regulated gene Rem2 as an important and novel regulator of distinct forms of homeostatic plasticity. Our findings demonstrate that Rem2 functions in excitatory cortical neurons to mediate OD plasticity during the critical period (Fig. 3 and 4). We show that in the absence of Rem2 both homeostatic postsynaptic scaling up (Fig. 5) and neuronal intrinsic excitability (Fig. 6) are altered. We further reveal that Rem2 is required cell-autonomously to establish the intrinsic excitability of layer 2/3 pyramidal neurons independent of changes in synaptic inputs (Fig. 7). Taken together with our previously published data (Ghiretti et al., 2013; Ghiretti et al., 2014; Ghiretti and Paradis, 2011), we propose that Rem2 functions as a transducer of Ca^2+^-dependent signals into long-lasting, functional changes in cell-intrinsic excitability and circuit connectivity. Importantly, our discovery of Rem2 as a key regulator of cortical plasticity provides a genetic inroad towards unraveling the signaling networks linking neural activity to functional output.

### Rem2 regulates visual system plasticity

Using the mouse visual system as a model of experience-dependent plasticity, we discovered that Rem2 is an important regulator of homeostatic plasticity mechanisms. Homeostatic processes work to stabilize circuit function in response to altered network activity through regulation of synaptic strengths, intrinsic excitability, and inhibition (Turrigiano, 2012). While several studies have examined the contributions of excitatory synaptic scaling up and cortical inhibition, little is known of the effect of cell-autonomous regulation of intrinsic excitability on visual circuit function. Our electrophysiological studies demonstrate that the activity-regulated gene *Rem2* is required for homeostatic synaptic scaling up (Fig. 5) and proper regulation of intrinsic excitability *in vivo* (Fig. 6). Importantly, our data reveal that Rem2 establishes the intrinsic excitability of neurons in an acute and cell-autonomous manner in the absence of any detectable change in excitatory synaptic inputs or sensory input (Fig. 7). Thus, the primary function of Rem2 signaling is to set the intrinsic excitability of these neurons. Additionally, we demonstrate that deletion of *Rem2* from cortical excitatory neurons (*Rem2*^*flx*/*flx*^; *EMX1*^*Cre*^), but not PV^+^ or VIP^+^ interneurons, results in a failure to undergo OD plasticity, similar to *Rem2*^-/-^ animals (Fig. 4). Taken together, the deficit in late-phase OD plasticity and normal adult OD plasticity observed in *Rem2*^-/-^ mice (Fig. 3), impaired synaptic scaling up (Fig. 5), and altered intrinsic excitability (Fig. 6) support the premise that Rem2 regulates distinct homeostatic plasticity mechanisms during the critical period.

Interestingly, transgenic mice harboring a deletion of TNFα also display a deficit in late-phase OD plasticity (Kaneko et al., 2008b) and normal adult OD plasticity (Ranson et al., 2012). In this study, the authors demonstrate that impaired homeostatic scaling up results in a failure to increase open eye responsiveness, causing the observed deficit in OD plasticity (Kaneko et al., 2008b). However, this study did not examine a possible role for TNFα in regulation of intrinsic excitability. In contrast, *Rem2*^-/-^ mice display both impaired synaptic scaling up (Fig. 5) and altered intrinsic excitability (Fig. 6), as well as a generalized increase of individual eye responses with 6d MD (Fig. 3C). These latter results are similar to those observed following binocular deprivation, which leads to increased cortical responsiveness to both eyes (Mrsic-Flogel 2007). While it may seem counterintuitive to observe defective synaptic scaling yet increased cortical responsiveness, it is the combination of impaired synaptic scaling and aberrant intrinsic excitability that lead to the observed alterations in circuit output. There is no precedence in the literature for how this combination of cellular phenotypes would influence cortical responsiveness as assayed by ISI. Thus, while it remains to be determined exactly how these two processes function synergistically to maintain proper circuit function this data provides novel insight into the importance of neuronal regulation of intrinsic excitability to sculpting network output.

**Figure 7.**
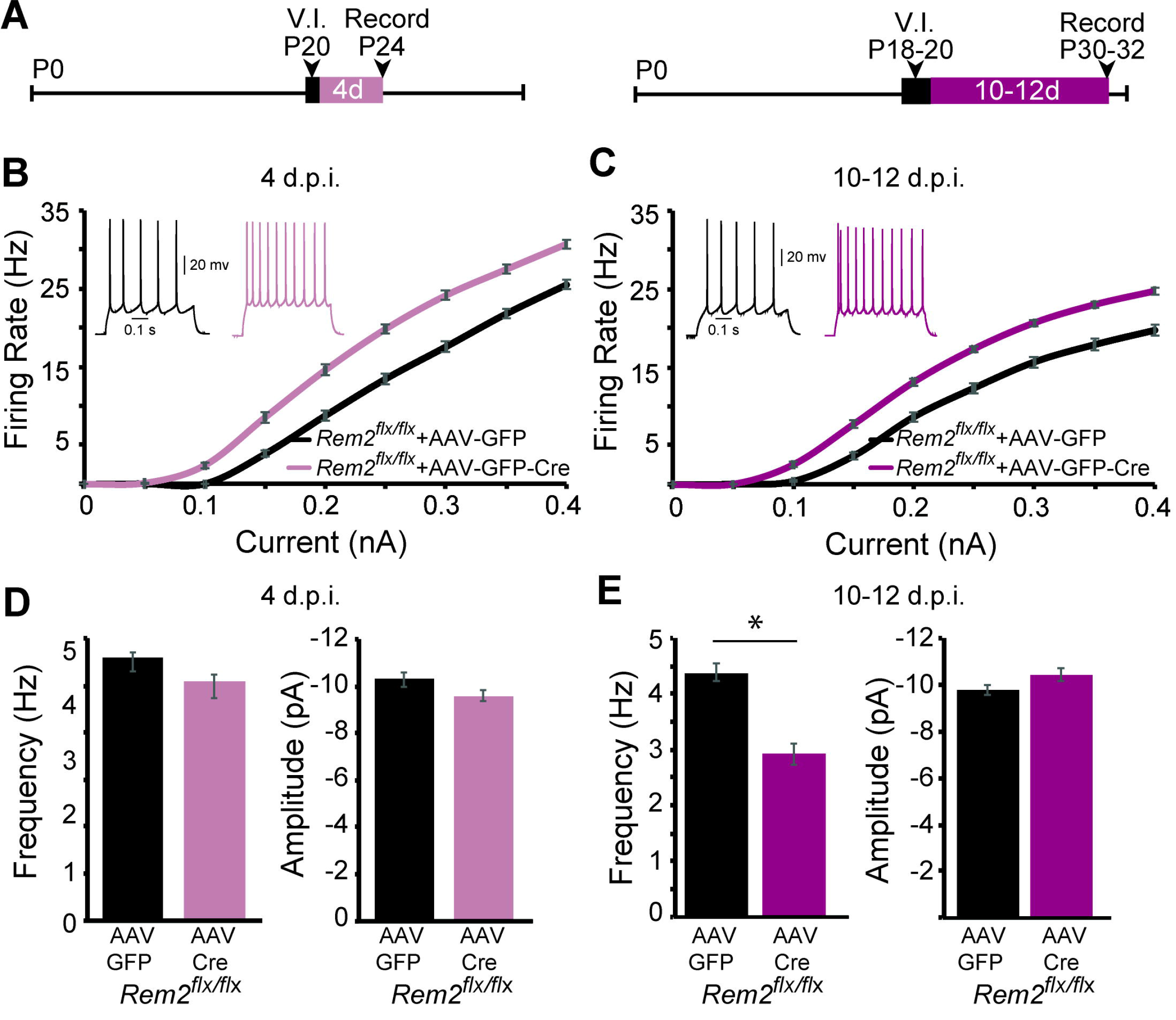
Rem2 cell-autonomously regulates intrinsic excitability *in vivo*. **A**) Experimental timeline of acute, viral-mediated *Rem2* deletion in *Rem2*^*flx*/*flx*^ mice. V.I., virus injection. **B**) Average *f-I* curves recorded from neurons of *Rem2*^*flx*/*flx*^ mice injected with either a control GFP (*Rem2*^*flx*/*flx*^ + AAV-GFP, black, n=17) or GFP-Cre expressing virus (*Rem2*^*flx*/*flx*^ + AAV-GFP-Cre, mauve, n=19) measured 4 days post infection (d.p.i.). Inset: representative traces of evoked responses measured at 0.2 nA from *Rem2*^*flx*/*flx*^ + AAV-GFP and *Rem2*^*flx*/*flx*^ + AAV-GFP-Cre neurons 4 d.p.i.․ **C**) Average *f-I* curves recorded from neurons of *Rem2*^*flx*/*flx*^ + AAV-GFP (black, n=18) or *Rem2*^*flx*/*flx*^ + AAV-GFP-Cre (magenta, n=20) measured 10-12 days post infection. Inset: representative traces of evoked responses measured at 0.2 nA from *Rem2*^*flx*/*flx*^ + AAV-GFP and *Rem2*^*flx*/*flx*^ + AAV-GFP-Cre neurons 10-12 d.p.i.․ **D**) Average mEPSC (left) frequency and (right) amplitude measured in *Rem2*^*flx*/*flx*^ + AAV-GFP (black, n=14) or *Rem2*^*flx*/*flx*^ + AAV-GFP-Cre (mauve, n=17) measured 4 days post infection. **E**) Average mEPSC (left) frequency and (right) amplitude measured in *Rem2*^*flx*/*flx*^ + AAV-GFP (black, n=22) or *Rem2*^*flx*/*flx*^ + AAV-GFP-Cre (magenta, n=18) measured 10-12 days post infection. n=3-5 animals per condition. Data is presented as mean ± SEM. *p < 0.05, by two-way ANOVA and Tukey post-hoc for mEPSC frequency and amplitude mean data plots.

It is interesting to note that the *Rem2*^*flx*/*flx*^; *EMX1*^*Cre*^ animals exhibit a smaller shift in ocular dominance after 6d MD than the *Rem2*^-/-^ animals when compared to their respective littermate controls (compare Fig. 3C to Fig. 4A). One possible explanation for this result is that Rem2 modulates plasticity via its expression in different cell types or brain regions other than those examined by use of individual Cre driver lines in the current study. Previous work investigating the role of *Igf1* in VIP^+^ interneurons in OD plasticity demonstrated that while deletion of *Igf1* in VIP^+^ interneurons altered inhibitory tone it did not affect OD plasticity following monocular deprivation (Mardinly et al., 2016). Thus, another possibility is that Rem2 may also function in other cell types, such as VIP^+^ interneurons, to help disinhibit cortical circuits and indirectly contribute to the partial shift observed in the *Rem2*^-/-^ animals that is absent in the *Rem2*^*flx*/*flx*^; *EMX1*^*Cre*^ mice. Taken together, our results are consistent with a role for Rem2 in mediating circuit function by regulating homeostatic mechanisms in excitatory, cortical neurons (e.g. postsynaptic scaling of excitatory synapses or changes in intrinsic excitability).

### Rem2 signaling in homeostatic plasticity

Our discovery of Rem2 function in homeostatic plasticity provides us with a unique entry point to discover the activity-dependent cytoplasmic signaling networks that regulate neuronal excitability. A number of activity-regulated genes have been implicated in both homeostatic synaptic scaling and OD plasticity including: Arc (Gao et al., 2010; McCurry et al., 2010; Shepherd et al., 2006), BDNF (Kaneko et al., 2008a; Rutherford et al., 1998), Homer1a (Hu et al., 2010; Nedivi, 1999), MHCI (Goddard et al., 2007; Syken et al., 2006), and NARP (Chang et al., 2010; Gu et al., 2013). Interestingly, all of these factors are either secreted, membrane-bound, or localized to and function at the postsynaptic density (Davis, 2013; Fernandes and Carvalho, 2016; Pozo and Goda, 2010). In addition, changes in gene expression underlie the long-lasting nature of homeostatic plasticity (Turrigiano, 2008). Accordingly, the activity-regulated transcription factor CREB as well as CaMKIV, a nuclear CaMK that activates CREB, have also been implicated in synaptic scaling (Benito and Barco, 2010; Ibata et al., 2008; Joseph and Turrigiano, 2017). Thus, up until our discovery that Rem2 regulates synaptic scaling, identification of activity-dependent, cytosolic signaling molecules that link changes at the neuronal membrane to changes in gene expression to regulate homeostatic plasticity were lacking.

While a number of molecules have been implicated in regulation of synaptic scaling, a comparable understanding of the regulation of neuronal intrinsic excitability is lacking (CREB (Dong et al., 2006) and CaMKIV (Joseph and Turrigiano, 2017)). Rem2 is a noncanonical Ras-like GTPase, expressed throughout the cell (Ghiretti and Paradis, 2011) and is unlikely to be regulated by its nucleotide binding state (Correll et al., 2008). The mechanism by which Rem2 transduces signals is an open area of investigation, although it has been shown to associate with VGCC subunits (Chen et al., 2005; Finlin et al., 2006; Finlin et al., 2005; Pang et al., 2010) and CaMKII (Flynn et al., 2012; Royer et al., 2017). In addition, our previous studies demonstrated that Rem2 functions in a CaMK signaling pathway to regulate dendritic complexity (Ghiretti et al., 2014). Interestingly, CREB is a known output of CaMK signaling and has also been implicated in regulation of dendritic morphology (Redmond et al., 2002), synaptic scaling (Ibata et al., 2008; Joseph and Turrigiano, 2017), and intrinsic excitability (Dong et al., 2006; Joseph and Turrigiano, 2017). In fact, two of these studies (Dong et al., 2006; Joseph and Turrigiano, 2017) strongly suggest that neuronal intrinsic excitability is under transcriptional control. We propose that Rem2 functions as a calcium-sensitive cytoplasmic signal transduction molecule, conveying changes at the membrane (e.g. MHCI signaling or Ca^2+^ influx) to changes in gene expression in the nucleus (e.g. CaMKIV, CREB) to regulate homeostatic plasticity.

How might Rem2 instruct changes in intrinsic excitability more specifically? Recent modeling studies illustrate that the rate of calcium-dependent gene transcription and channel conductance as one point of regulation underlying changes in intrinsic excitability and neuronal conductances (O'Leary et al., 2014). Interestingly, this model posits the existence of a key regulator or “sensor” molecule whose main function is to assess cell-wide activity levels, indicated by changes in Ca^2+^ influx, and implement downstream signaling mechanisms (Liu et al., 1998; O'Leary et al., 2014; Siegel et al., 1994). Given that Rem2 expression and signaling is sensitive to neuronal activity levels, and that a major output of Rem2 function is to regulate intrinsic excitability, we posit that Rem2 may be one such cytosolic sensor molecule. In support of this hypothesis, we have recently performed RNA-sequencing to identify downstream targets of Rem2 (Kenny et al., 2017). We found that the expression of a number of ion channels, important for establishing neuronal excitability, are regulated by Rem2 signaling in an activity-dependent manner (Kenny et al., 2017). Thus, Rem2 signaling may regulate neuronal intrinsic excitability by controlling the composition of ion channels at the cell membrane.

### Rem2 in structural plasticity

Our previous findings revealed that Rem2 is also an important regulator of synapse formation and dendritic morphology in cultured neurons and *Xenopus* optic tectum (Ghiretti et al., 2013; Ghiretti et al.․ 2014; Ghiretti and Paradis, 2011; Moore et al., 2013). The current study also demonstrates that Rem2 regulates dendritic spine density in response to sensory experience (Fig. 5D, S3). Similar to our findings with Rem2, other activity signaling pathways also co-regulate experience-dependent plasticity and synapse development (Piochon et al., 2016; Titley et al., 2017). For example, *cpg15* has been shown to be an important regulator of synapse formation and dendritic complexity and is required during the critical period for normal OD plasticity (Picard et al., 2014). Conversely, MHCI negatively regulates synapse development and deletion of MHCI or PirB, a MHCI receptor, enhances visual system plasticity *in vivo* (Cebrian et al., 2014; Shatz, 2009). Although the exact contribution of subtle changes in excitatory synapse density to visual circuit function in general and OD plasticity in particular is not well understood, Hofer et al., demonstrated that spine formation and turnover is dramatically affected by MD in layer 5 cortical neurons (Hofer et al., 2009), suggesting that an important functional correlation exists. Hence, our findings that Rem2 functions as a regulator of experience-dependent plasticity at the morphological, cellular and circuit levels provide an important step forward in connecting molecular regulators of neuronal morphology with broader circuit function.

### Conclusions

In conclusion, our *in vivo* analysis of Rem2 function in the visual system unveils a unique role for Rem2 as a key regulator of neural circuit plasticity mechanisms. In the future, defining the molecular mechanisms that underlie Rem2-dependent homeostatic plasticity, specifically cell-autonomous regulation of intrinsic excitability, will elucidate the genetically encoded networks that instruct activity-dependent modifications of neural circuitry. In addition to cortex, Rem2 is expressed in other areas of the brain including the hippocampus, amygdala, and nucleus accumbens (Ghiretti and Paradis, 2011; Liput et al., 2016), which are critical regions for learning, memory, and addiction. Interestingly, these processes rely heavily on physiological and morphological plasticity at the level of individual neurons (El-Gaby et al., 2015; Luscher and Malenka, 2011; Martin et al., 2000; Yager et al., 2015). Therefore it is likely that Rem2 acts in other brain regions and in adulthood, based on transcriptional profiling experiments performed in adult animals (Liput et al., 2016; Mardinly et al., 2016). Our data indicate that Rem2 functions at the nexus of a signaling network that senses and responds to changes in neuronal activity in order to preserve proper circuit function in the face of changing sensory experience.

## EXPERIMENTAL PROCEDURES

All experimental procedures involving animals were approved by the Institutional Animal Care and Use Committee at Brandeis University.

### Western blot

Rat or mouse cortices were isolated at different developmental ages (see Fig. 1A) and homogenized on ice in RIPA buffer + 1x Complete Protease Inhibitor Tab (Roche) using 21, 23 and 30 gage needles attached to a 3ml syringe. Total protein concentration for each lysate was determined using a Bio-Rad protein assay and equal amounts were loaded on the gel for each condition. Lysates were mixed with homemade 3x sample buffer (6% SDS, 0.1% bromophenol blue, 150 mM Tris, pH 6.8, 30% glycerol, and 10% β-mercaptoethanol) and boiled for 7 minutes. The lysates were run on a 12% SDS-PAGE gel and the proteins were transferred to a nitrocellulose membrane. The membrane was probed using anti-Rem2 (1:500; Santa Cruz cat # sc160722; RRID: AB_2179340) and anti-βactin (1:5000; Abcam cat # ab 8226; RRID: AB_306371) antibodies as a loading control. Western blots were developed using the Odyssey Infrared Imaging System (Licor).

### Visual Stimulation and Gene Expression Analysis

Young mice were reared from postnatal day 9 (P9, prior to eye opening) until P28 in a light-tight dark box (Phenome Technologies, Inc). At P28, one group was left in the dark while a second group was exposed to light for 90 min. Then mice were anesthetized with isoflurane and decapitated. Mice from the dark group were decapitated and brains removed with the use of night vision goggles. The visual and somatosensory cortices were isolated using a customized tissue punch based on age-matched anatomical reference points and RNA was extracted with Trizol Reagent (Invitrogen). DNase-free RNA was prepared and then reverse transcribed using a Random Primer Mix (New England Biolabs). Quantitative real-time PCR was performed using SYBR green detection (Clontech) on a Rotor Gene thermal cycler (Roche). Primers for *Rem2* and *Fos* were previously verified in Ghiretti et al., 2014. The PCR products were normalized to *Actb* (β-actin) and presented as fold change over baseline using the ΔΔCT method. “n” represents the number of biological replicates used. Data were compiled from independent experiments each conducted in triplicate. A one-way ANOVA followed by a Dunnett's test was used to compare experimental conditions to control or stimulation.

### Generation of Rem2^−/−^ and Rem2^flx/flx^ mice

Embryonic stem cell lines harboring a reporter-tagged insertion with conditional potential at the *Rem2* locus (referred to as *Rem2*^-/-^; Fig 2A) were acquired from the European Conditional Mouse Mutagenesis Program (EUCOMM) from the International Knockout Mouse Consortium (IKMC; Ref ID: 92501). The cassette was inserted into the first intron of the *Rem2* locus and contained a mouse *En2* splice acceptor sequence (EN2), IRES, a LacZ gene, a SV40 polyadenylation, and a Neo gene flanked by FRT sites and LoxP sites flanking exons 2 and 3 (see Fig. 2A; see Skarnes et al., 2011 for details). Insertion of the cassette into the *Rem2* locus in the ES lines was verified by extensive PCR and sequencing. Embryonic stem cells were injected into C57BL/6 blastocysts using standard conditions. Injected blastocysts were surgically implanted into pseudo-pregnant foster female mice to generate chimeric offspring. Chimeras were mated to C57BL/6 females to obtain germline transmission; genotyping was performed by PCR. Correct insertion of the cassette was verified by Southern blotting (see Fig. 2B). Note that for unknown reasons, β-galactosidase protein was not expressed in mice harboring the *Rem2* null allele. For routine experimentation, animals were genotyped using a PCR-based strategy. Animals harboring the *Rem2* null allele were genotyped with a forward primer in the intron spanning region of exon 1 (5’-GCTTCTTCTAGCTCCATCGTTG-3’) and a reverse primer in the inserted cassette region (5’-GGACCACCTCATCAGAAGC-3’) or the reverse primer in exon 2 (5’-AGTTGGGAAGCTATATCTTC-3’) (Fig. 2A, blue arrows, D).

We observed that inbreeding of the *Rem2*^-/-^ allele to the C75BL/6 strain resulted in mice with small litters and poor viability. To determine if this phenotype was due to deletion of the *Rem2* gene, we outcrossed these mice to the 129-Elite mice strain (Charles River) and found a dramatic improvement in viability and litter size. Thus we continued to outcross the *Rem2*^-/-^ allele for at least 5 more generations. All experiments reported in this study for the *Rem2*^-/-^ allele are on this outcrossed background. We used aged-matched, littermate WT and *Rem2*^-/-^ mice for all experiments. Note that in a handful of experiments for intrinsic signal imaging, a complete set of age-matched littermate controls were not always possible (see below).

To generate a *Rem2* conditional allele, the *Rem2*^-/-^ mice were crossed with mice expressing Flp recombinase with a ROSA26 promoter region (JAX 009086) resulting in exons 2 and 3 flanked by loxP sites (Fig. 2A). Conditional knockout animals were genotyped for the presence of the remaining loxP site, which shifts the size of the PCR product in the *Rem2*^*flx*/*flx*^ animal by 66 bp, with the forward primer (5’-CATCCTGGCTCCAACCATGG-3’) and reverse primer (5’-CTCCGGTCCTGTCACATCAG-3’) in the intron spanning region between exons 3 and 4 (Fig. 2D). These mice were also backcrossed into the 129-Elite mice for more than 5 generations. To generate lines with *Rem2* deleted in defined cell types, *Rem2*^*flx*/*flx*^ mice were crossed to either the EMX1Cre line (*EMX1*^*Cre*^, JAX #005628) or PVCre line (*PV*^*Cre*^, JAX #017320). All mice were maintained as heterozygous mating pairs. Mutant mice were identified by performing PCR on tail genomic DNA (Fig. 2D).

### Southern blot

Genomic DNA was isolated from the livers of mice and digested with Sca1 and Asc1 restriction enzymes. The DNA was run on a 0.8% agarose gel and transferred to a nitrocellulose membrane using a vacuum blotter (Qbiogene, TransDNA Express™ Vacuum Blotter). The nitrocellulose membrane was incubated in pre-hybridization solution (50% Formamide, 5x SSPE, .1% SDS, 5x Denhardt's solution, 1mg Salmon sperm DNA) at 42°C for 4 hours. The probes were labeled using α-^32^P dATP (3000Ci/mmol; PerkinElmer Life and Analytical Sciences) and the Prime-it II Random Primer labeling kit (Stratagene) according to manufacturer's instructions. The probes were added to hybridization buffer (50% Formamide, 5x SSPE, .1% SDS, 1x Denhardt’s solution, 1mg Salmon sperm DNA) and incubated overnight at 42°C. The nitrocellulose was washed three times with 2x SSC/0.1% SDS, once with 0.5x SSC/0.1% SDS and once with 0.1x SSC/0.1% SDS. The membrane was exposed overnight in a phosphorlmager cassette and imaged using the STORM molecular imaging system (GE Healthcare). The inserted 7.5kb cassette introduced an AscI site between the LacZ and Neo markers (Fig. 2A). Therefore, the predicted band size with the 3’ probe, if the cassette was correctly inserted at the *Rem2* locus, is an 11 kb band for the *Rem2*^-/-^ locus and a 13.8 kb band for the wildtype locus (Fig. 2B) using a ScaI/AscI double digest.

### Cortical Thickness and Brain Weight Measurements

Typically reared P7, P21, and P30 WT or *Rem2*^-/-^ littermates were deeply anesthetized using ketamine/xylazine cocktail (ketamine 50mg/kg, xylazine 5mg/kg) and perfused first with 0.1M PBS followed by 4% paraformaldehyde in 0.1M sodium phosphate buffer. Brains were carefully extracted and stored in 4% paraformaldehyde in 0.1M sodium phosphate buffer for 24 hours, then transferred to 30% sucrose for at least 24 hours. Sections were cut at 30 μm on a freezing microtome and mounted on slides coated with 2% porcine gelatin. Slides were allowed to dry at room temperature and stored at 4°C until histology was performed. Briefly, slides were washed with xylenes and then dehydrated with a series of graded ethanols (100%, 95%, and 70%). Slides were then rinsed with ddH_2_0, stained with 0.1% cresyl violet for 1 minute, cleared with a grades series of ethanols, and differentiated with glacial acetic acid in 95% ethanol. Slides were then dehydrated in a series of graded ethanols, cleared with xylenes, and coverslipped using Permount (Fisher Scientific). Every second section containing visual cortex was imaged at 4X magnification using a Keyence BZX-700 microscope (Keyence). Visual cortex was identified using anatomical landmarks diagramed in the Allen Brain Atlas and cortical layers were identified using changes in cell size and density. Cortical thickness was measured from the deepest extent of layer 6 to the cortical surface using the ImageJ Measure function. Boundaries of cortical layers were manually drawn using ImageJ. Averages per animal are computed across all measured sections of visual cortex. Resulting measurements from *Rem2*^-/-^ mice were normalized to the measurements of their WT littermate to account for differences in histology conditions. Brain to body weight ratio was calculated using the weight and brain weight measurements recorded at the time of perfusion.

### Golgi-Cox Labeling

Typically reared WT and *Rem2*^-/-^ littermate mice were housed in a 12hr light/12hr dark cycle from birth to P30. Dark reared mice were placed in a light-tight box beginning at P9 until termination of the experiment (P30). At the specified age, the mice were anesthetized with ketamine/xylazine cocktail (ketamine 50mg/kg, xylazine 5mg/kg) and transcardially perfused with 0.9% saline in ddH_2_0. Dark-reared mice were anesthetized in the dark and then shielded from light until after perfusion using light blocking tape (Thor Labs) to cover the eyes. Immediately following perfusion, brains were weighed and submerged in Golgi-Cox solution (FD NeuroTechnologies). Throughout all steps involving Golgi-Cox, brains were protected from light. Golgi-Cox solution was changed 24 hours after initial immersion and brains continued to be stored in Golgi-Cox solution for 7 days. After 7 days, brains were transferred to Solution C (FD Neurotechnologies) for at least 2 days. Sections were cut at 150 μm using a cryostat and immediately mounted on slides coated with 2% porcine gelatin. Histology was carried out according to the protocol supplied by FD Neurotechnologies RapidGolgi Stain Kit. Briefly, slides were washed with ddH_2_O, developed using FD Neurotechnologies Solutions D & E, rinsed in ddH_2_O, dehydrated with a graded series of ethanols, and cleared using xylenes. Slides were then coverslipped using Permount (Fisher Scientific).

Tissue sections were imaged in brightfield using a Zeiss AxioObserver microscope. For spine density quantification, z-stacks of images were captured using a 60X oil objective. Neurons were sampled from layer 2/3 pyramidal neurons of the visual cortex. Visual cortex was identified using anatomical landmarks diagramed in the Allen Brain Atlas and pyramidal neurons were identified by their well-described somatic and dendritic morphology. Entire terminal apical branches approximately 50 - 100 μm from the soma were chosen for reconstruction and spine density quantification based on absence of artifact, lack of structural damage, and completeness of staining. Image stacks include the entire branch beginning at the branch point with the apical trunk and continuing to the branch tip. For cortical thickness measurements, single images of each section throughout the anterior-to-posterior extent of visual cortex were captured using brightfield illumination at 4X magnification using a Keyence BZX-700 microscope. All analysis was performed with the experimenter blind to genotype and rearing condition. Dendritic spines were counted manually in FIJI (NIH) on z-stack images using the Cell Counter plugin. A dendritic spine was identified as any protrusion from the dendritic segment at least 0.5 μm in length or greater. Dendritic segment length was measured using the NeuronJ plugin. A total of 1500 μm of dendritic segment was counted for the WT and *Rem2*^-/-^ TR segments and a total of 2000 μm of dendritic segment was quantified for the WT and *Rem2*^-/-^ DR images, for an average of 50 μm segment measured per neuron.

### Calcium Imaging

Dissociated cortical neurons from embryonic day 16 (E16) mice from a *Rem2*^+/−^ x *Rem2*^+/−^ mating were plated on 12mm glass coverslips at a density of approximately 70,000/cm^2^ and grown in glia-conditioned Neurobasal media with B27 supplement (Invitrogen). E16 littermates were dissociated side-by-side and plated on individual coverslips and subsequently genotyped by PCR to determine whether the cultured neurons were *Rem2*^+/+^, Rem2^+/−^, or *Rem2*^-/-^. Free resting calcium measurements were conducted 5 days after plating (DIV 5). Cortical neurons were incubated with 1 μM Fura2-AM (Invitrogen) in Tyrode's solution containing 0.1% bovine serum albumin for 30 min at 37°C. The neuronal culture was then washed with Tyrode's solution and incubated for an additional 30 min at 37°C in the same solution for de-esterification. After the incubation period, the coverslip was washed once more with Tyrode's solution and mounted on an imaging chamber containing the same solution. Fura-2 fluorescence images were acquired at 37°C on an Olympus IX-70 inverted microscope using a 20x 0.7 NA objective (Olympus UPlanApo) and a cooled CCD camera (Orca R2, Hamamatsu) controlled by Volocity software (Improvision). Fluorophore excitation was achieved using a mercury lamp and spectral separation for excitation and emission was obtained using a fura-2 filter set (Brightline Fura2-C, Semrock). Fura-2 fluorescence images with excitation centered at 340 nm and 387 (26 and 11 nm bandwidth, respectively) and emission collected at 468-552 nm were acquired sequentially using a motorized filter-wheel (Prior Scientific).

### Lid Suturing

Wildtype or *Rem2*^-/-^ littermates were either not surgerized or monocularly lid sutured between P25-P27 for a duration of either 2 days (2d MD) or 5-7 days (6d MD). Mice were anesthetized using either a ketamine/xylazine cocktail (ketamine 50mg/kg, xylazine 5mg/kg) or 2.0% isoflurane delivered using a SomnoSuite vaporizer system (Kent Scientific). The surgical area was cleaned thoroughly using povidone pads (Dynarex), with care taken to avoid contact with the eye. The eye to be suture was rinsed using bacteriostatic saline and coated with a thin film of antibacterial ophthalmic ointment prior to suturing. Lid margins were trimmed and the lid was closed with 2-3 mattress sutures using silk suture thread. Additional antibacterial ointment was applied to the sutured lid. Sutures were checked daily and if not intact animals were not used.

Lid suture for adult OD plasticity experiments was completed as above, but with suturing occurring instead between the ages of P60-74.

### Intrinsic Signal Imaging

Mice aged between P31 and P33 underwent intrinsic signal imaging (ISI) for critical period OD plasticity experiments and adult mice (10-12 weeds of age) underwent ISI for adult OD plasticity experiments. Experimental did not differ between juvenile and adult groups. The experimenter was blind to genotype until the conclusion of the experiment and data analysis. The genotype of each mouse was confirmed postmortem. Mice were prepared for ISI experiments in littermate cohorts such that littermate controls were used for both experimental factors (deprivation and genotype). In the event that a littermate was not usable, such as an open eyelid suture or death during an experiment, the remaining littermate was included in our data. Analysis of our results with these mice removed resulted in the same significant relationships between groups that we report in the manuscript.

Anesthesia was induced using 4% isoflurane (100mL/min) in an anesthesia chamber delivered using a SomnoSuite (Kent Scientific) and anesthesia was maintained at 2% isoflurane during surgical procedures. Isoflurane anesthesia was supplemented with a single dose of chlorprothixene (10mg/kg). First, mice that had previously undergone lid suture had sutures removed and the eye re-opened. Each sutured eye was closely inspected for health and clarity; any mice with clouding of the eye or infection of the surgical area were euthanized.

In preparation for ISI, an incision was made on the scalp and skin resected to expose the skull. The skull was cleared of overlying tissue and dried to allow for secure attachment of a small headpost using adhesive (ZAP Gel, Pacer Technologies). Following placement of the headpost, mice were headfixed and the skull was thinned over a wide area containing visual cortex contralateral to the deprived eye (if applicable) using a scalpel (blade 15) until transparent under saline. The location of visual cortex was estimated as 3.0mm lateral to the midline and 1mm anterior to the lambda suture. After thinning, an optical widow was created to stabilize the skull and provide a flat imaging surface by applying 2% agarose to the skull and pressing a 5 mm coverslip into the agarose. This window was sealed and affixed to the skull using cyanoacrylate adhesive.

Following surgical procedures, anesthesia was reduced to between 0.7% and 1.0% for imaging. Anesthesia level was kept as low as possible without leading to pain response in the mouse. Before ISI, the brain was imaged using green light (575 nm) to serve as a record of the imaging field's clarity and health. Intrinsic signal imaging was performed under red light (675 nm) and images were captured at a rate of 30Hz by a Dalsa 21−01M60 camera. Mice were shown 20 repetitions of drifting grating stimuli at either 0° or 90° of 100% contrast at 2Hz temporal frequency and 0.04cycles/degree spatial frequency lasting for 10 seconds with an interstimulus interval of 15 seconds. A blank control screen of equal average luminance to the drifting gratings was also presented as a control stimulus. Stimuli were presented with stimuli to each eye alternately, with visual input being blocked to the opposing eye using opaque, flexible light-blocking material.

All image analysis was performed using custom Matlab (Mathworks) scripts written in the Van Hooser Lab. Acquired images were mean filtered, blank screen subtracted, and summed across all frames. A region of interest over binocular visual cortex, the area of the cortex responding to stimulation of the ipsilateral eye, was manually drawn. An ocular dominance index (ODI) was calculated as follows, were R_C_ is the response to stimulation of the eye contralateral to the imaging window and R_I_ is the response to stimulation of the eye ipsilateral to the imaging window.

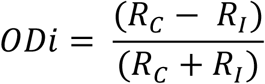

### Virus Injection

The AAV-GFP (AAV1.hSyn.eGFP.WPRE.bGH; AV-1-PV1696) and AAV-GFP-Cre (AAV1.hSyn.HI.eGFP-Cre.WPRE.SV40; AV-1-PV1848) constructs were obtained from the Vector Core facility at the University of Pennsylvania. Mice age P18-P20 were anesthetized with a cocktail of ketamine (100 mg/kg) and xylazine (10mg/kg), mounted on a stereotaxic frame, and the scull exposed. Primary binocular visual cortex was targeted by using the mouse brain atlas after adjusting for the lambda-bregma distance for age. A small 1-2 mm in diameter hole was drilled in the skull and a glass micropipette delivered 50 nL AAV (diluted 1:250) 150μm below the dural surface. The scalp was sutured and betadine was applied. Animals recovered on a heating pad and were returned to the animal facility until use.

### Electrophysiology

Whole-cell patch clamp recordings were performed on layer 2/3 pyramidal neurons in binocular primary visual cortex (V1b). Mice were anesthetized with isoflurane, decapitated, and the brains removed with the head immersed in ice-cold, choline-based cutting solution. The choline cutting solution contained (mM) 25 NaHCO_3_, 1.25 NaH_2_PO_4_·H_2_O, 2.5 KCl, MgCl_2_·6H_2_O, 25 glucose, 0.5 CaCl_2_, 110 C_5_H_14_ClNO, 11.6 ascorbic acid, and 3.1 pyruvic acid. Coronal slices (300 μm) were cut on a vibrating microtome (Leica) and allowed to recover at 37°C for 29 minutes followed by an additional 29 minutes at room temperature before recording. V1b was identified using the mouse brain atlas and the anatomical landmarks of the shape and morphology of the white matter (Lambo and Turrigiano 2013; Maffei and Turrigiano 2008). Layer 2/3 pyramidal neurons located in the center of V1b were selected for recordings to avoid boundary regions identified with a 40x immersion lens. Recordings were obtained at 34°C with a Multiclamp 700B amplifier and a Digidata 1440A digitizer controlled by pClamp10 software. R_s_ and R_IN_ were monitored throughout experiments using Lab Bench (Clampex 10.2) and cells that exhibited changes of more than 20% in either of these parameters throughout the course of the recording were discarded. For AMPA-mediated mEPSC recordings the external solution contained (in mM): 125 NaCl, 26 NaHCO_3_, 2.3 KCl, 1.26 KH_2_PO_4_, 2 CaCl_2_, 2 MgSO_4_, 10 glucose, 1 μM tetrodotoxin (TTX, Abcam Biochemicals), 50 μM DL-2-Amino-5-phosphonopentanoic acid (APV; Sigma Aldrich), and 50 μM Picrotoxin (Sigma). The internal pipette solution contained (in mM): 135 Potassium gluconate, 10 HEPES, 2 MgCl2, 3 Na_2_ATP, 0.3 Na_2_GTP, and 10 Phosphocreatine (pH 7.3, adjusted with KOH). Rhodamine was included in the recording pipette to verify neuronal morphology at the conclusion of the recording period. The membrane potential was held at −70 mV and events were filtered at 1 kHz. Data was recorded in 3 epochs at 100 s each for a total duration of 300s per cell. mEPSCs were then evaluated offline in Clampfit 10.2 (Molecular Devices). Detection criteria of mEPSC events included amplitudes > 5 pA and rise times < 3 ms. To measure action potential firing rates (*f-I* curves), a series of step pulses (duration 600 ms) from −200 to 400 pA in 50 pA increments were delivered from rest in the presence of 50 μM APV, 25 μM DNQX, and 50 μM Picrotoxin. A small bias current was injected to maintain V_m_ at −70 mV in between depolarizations.

Measures of mIPSCs were done in the presence of 1 μM TTX, 50 μM APV, and 10 μM DNQX (Sigma Aldrich) to isolate inhibitory postsynaptic currents. Patch pipettes (3 - 5 MΩ) were filled with intracellular solution containing (in mM) 120 CsCl, 10 HEPES, 1 EGTA, 0.1 CaCl_2_, 1.5 MgCl_2_, 4 Na_2_ATP, and 0.3 Na_2_GTP (pH 7.3, adjusted with CsOH). Data is presented as mean ± S.E.M. Statistical significance was calculated using a two-way ANOVA and Tukey post hoc analysis for averaged data. Kolmogorov-Smirnov test was used to test for comparison in the cumulative distribution plots.

## Author Contributions

Conceptualization A.R.M, S.E.R., S.D.V., S.P.; Methodology A.R.M., S.E.R., K.K., L.R., U.C., and K.F.; Writing - Original Draft, A.R.M. and S.P.; Writing - Review and Editing, S.E.R., K.K., L.R., U.C., K.F., and S.D.V.; Funding Acquisition, A.R.M., S.D.V., and S.P.; Supervision, S.D.V. and S.P.

## Acknowledgments

We thank Drs. Eve Marder, Gina Turrigiano, and Amy Ghiretti for advice and comments on the manuscript, the Paradis and Van Hooser labs for helpful comments and suggestion throughout the project and Lyra Hall and Katherine Kimbrell for technical assistance. This work was supported by the Charles Hood Foundation (S.D.V.), the Ruth L. Kirschstein NIH Training Grant T32NS007292 (A.R.M.), the NIH Career Development K01 award K01MH101639 (A.R.M.), NIH grant EY022122 (S.D.V.) and R01NS065856 (S.P). We would like to thank Dr. Margaret Thompson and acknowledge the IDDRC Mouse Gene Manipulation Core (NIHP30-HD 18655) for their help with generation of the *Rem2*^-/-^ mice.

## REFERENCES

Aizenman, C.D., Akerman, C.J., Jensen, K.R., and Cline, H.T. (2003). Visually driven regulation of intrinsic neuronal excitability improves stimulus detection in vivo. Neuron 39, 831–842.

Benito, E., and Barco, A. (2010). CREB's control of intrinsic and synaptic plasticity: implications for CREB-dependent memory models. Trends Neurosci 33, 230–240.

Butt, S.J., Fuccillo, M., Nery, S., Noctor, S., Kriegstein, A., Corbin, J.G., and Fishell, G. (2005). The temporal and spatial origins of cortical interneurons predict their physiological subtype. Neuron 48, 591–604.

Cebrian, C., Loike, J.D., and Sulzer, D. (2014). Neuronal MHC-I expression and its implications in synaptic function, axonal regeneration and Parkinson's and other brain diseases. Front Neuroanat 8, 114.

Chang, M.C., Park, J.M., Pelkey, K.A., Grabenstatter, H.L., Xu, D., Linden, D.J., Sutula, T.P., McBain, C.J., and Worley, P.F. (2010). Narp regulates homeostatic scaling of excitatory synapses on parvalbumin-expressing interneurons. Nat Neurosci 13, 1090–1097.

Chen, H., Puhl, H.L., 3rd, Niu, S.L., Mitchell, D.C., and Ikeda, S.R. (2005). Expression of Rem2, an RGK family small GTPase, reduces N-type calcium current without affecting channel surface density. J Neurosci 25, 9762–9772.

Cooke, S.F., and Bear, M.F. (2010). Visual experience induces long-term potentiation in the primary visual cortex. J Neurosci 30, 16304–16313.

Cooke, S.F., and Bear, M.F. (2014). How the mechanisms of long-term synaptic potentiation and depression serve experience-dependent plasticity in primary visual cortex. Philos Trans R Soc Lond B Biol Sci 369, 20130284.

Correll, R.N., Pang, C., Niedowicz, D.M., Finlin, B.S., and Andres, D.A. (2008). The RGK family of GTP-binding proteins: regulators of voltage-dependent calcium channels and cytoskeleton remodeling. Cell Signal 20, 292–300.

Crozier, R.A., Wang, Y., Liu, C.H., and Bear, M.F. (2007). Deprivation-induced synaptic depression by distinct mechanisms in different layers of mouse visual cortex. Proc Natl Acad Sci U S A 104, 1383–1388.

Davis, G.W. (2013). Homeostatic signaling and the stabilization of neural function. Neuron 80, 718–728.

Desai, N.S., Cudmore, R.H., Nelson, S.B., and Turrigiano, G.G. (2002). Critical periods for experience-dependent synaptic scaling in visual cortex. Nat Neurosci 5, 783–789.

Desai, N.S., Rutherford, L.C., and Turrigiano, G.G. (1999). Plasticity in the intrinsic excitability of cortical pyramidal neurons. Nat Neurosci 2, 515–520.

Dong, Y., Green, T., Saal, D., Marie, H., Neve, R., Nestler, E.J., and Malenka, R.C. (2006). CREB modulates excitability of nucleus accumbens neurons. Nat Neurosci 9, 475–477.

El-Gaby, M., Shipton, O.A., and Paulsen, O. (2015). Synaptic Plasticity and Memory: New Insights from Hippocampal Left-Right Asymmetries. Neuroscientist 21, 490–502.

Fernandes, D., and Carvalho, A.L. (2016). Mechanisms of homeostatic plasticity in the excitatory synapse. J Neurochem.

Fiala, J.C. (2005). Reconstruct: a free editor for serial section microscopy. J Microsc 218, 52–61.

Finlin, B.S., Correll, R.N., Pang, C., Crump, S.M., Satin, J., and Andres, D.A. (2006). Analysis of the complex between Ca2+ channel beta-subunit and the Rem GTPase. J Biol Chem 281, 23557–23566.

Finlin, B.S., Mosley, A.L., Crump, S.M., Correll, R.N., Ozcan, S., Satin, J., and Andres, D.A. (2005). Regulation of L-type Ca2+ channel activity and insulin secretion by the Rem2 GTPase. J Biol Chem 280, 41864–41871.

Finlin, B.S., Shao, H., Kadono-Okuda, K., Guo, N., and Andres, D.A. (2000). Rem2, a new member of the Rem/Rad/Gem/Kir family of Ras-related GTPases. Biochem J 347 Pt 1, 223–231.

Flynn, R., Labrie-Dion, E., Bernier, N., Colicos, M.A., De Koninck, P., and Zamponi, G.W. (2012). Activity-dependent subcellular cotrafficking of the small GTPase Rem2 and Ca2+/CaM-dependent protein kinase IIalpha. PLoS One 7, e41185.

Frenkel, M.Y., and Bear, M.F. (2004). How monocular deprivation shifts ocular dominance in visual cortex of young mice. Neuron 44, 917–923.

Fu, Y., Kaneko, M., Tang, Y., Alvarez-Buylla, A., and Stryker, M.P. (2015). A cortical disinhibitory circuit for enhancing adult plasticity. Elife 4, e05558.

Fu, Y., Tucciarone, J.M., Espinosa, J.S., Sheng, N., Darcy, D.P., Nicoll, R.A., Huang, Z.J., and Stryker, M.P. (2014). A cortical circuit for gain control by behavioral state. Cell 156, 1139–1152.

Gao, M., Sossa, K., Song, L., Errington, L., Cummings, L., Hwang, H., Kuhl, D., Worley, P., and Lee, H. K. (2010). A specific requirement of Arc/Arg3.1 for visual experience-induced homeostatic synaptic plasticity in mouse primary visual cortex. J Neurosci 30, 7168–7178.

Ghiretti, A.E., Kenny, K., Marr, M.T., 2nd, and Paradis, S. (2013). CaMKII-dependent phosphorylation of the GTPase Rem2 is required to restrict dendritic complexity. J Neurosci 33, 6504–6515.

Ghiretti, A.E., Moore, A.R., Brenner, R.G., Chen, L.F., West, A.E., Lau, N.C., Van Hooser, S.D., and Paradis, S. (2014). Rem2 is an activity-dependent negative regulator of dendritic complexity in vivo. J Neurosci 34, 392–407.

Ghiretti, A.E., and Paradis, S. (2011). The GTPase Rem2 regulates synapse development and dendritic morphology. Dev Neurobiol 71, 374–389.

Goddard, C.A., Butts, D.A., and Shatz, C.J. (2007). Regulation of CNS synapses by neuronal MHC class I. Proc Natl Acad Sci U S A 104, 6828–6833.

Gordon, J.A., and Stryker, M.P. (1996). Experience-dependent plasticity of binocular responses in the primary visual cortex of the mouse. J Neurosci 16, 3274–3286.

Gorski, J.A., Talley, T., Qiu, M., Puelles, L., Rubenstein, J.L., and Jones, K.R. (2002). Cortical excitatory neurons and glia, but not GABAergic neurons, are produced in the Emx1-expressing lineage. J Neurosci 22, 6309–6314.

Gu, Y., Huang, S., Chang, M.C., Worley, P., Kirkwood, A., and Quinlan, E.M. (2013). Obligatory role for the immediate early gene NARP in critical period plasticity. Neuron 79, 335–346.

Heimel, J.A., Hartman, R.J., Hermans, J.M., and Levelt, C.N. (2007). Screening mouse vision with intrinsic signal optical imaging. Eur J Neurosci 25, 795–804.

Hensch, T.K. (2005). Critical period plasticity in local cortical circuits. Nat Rev Neurosci 6, 877–888.

Hensch, T.K., and Fagiolini, M. (2005). Excitatory-inhibitory balance and critical period plasticity in developing visual cortex. Prog Brain Res 147, 115–124.

Hofer, S.B., Mrsic-Flogel, T.D., Bonhoeffer, T., and Hubener, M. (2006). Prior experience enhances plasticity in adult visual cortex. Nat Neurosci 9, 127–132.

Hofer, S.B., Mrsic-Flogel, T.D., Bonhoeffer, T., and Hubener, M. (2009). Experience leaves a lasting structural trace in cortical circuits. Nature 457, 313–317.

Hu, J.H., Park, J.M., Park, S., Xiao, B., Dehoff, M.H., Kim, S., Hayashi, T., Schwarz, M.K., Huganir, R.L., Seeburg, P.H., et al. (2010). Homeostatic scaling requires group I mGluR activation mediated by Homer1a. Neuron 68, 1128–1142.

Ibata, K., Sun, Q., and Turrigiano, G.G. (2008). Rapid synaptic scaling induced by changes in postsynaptic firing. Neuron 57, 819–826.

Joseph, A., and Turrigiano, G.G. (2017). All for One But Not One for All: Excitatory Synaptic Scaling and Intrinsic Excitability Are Coregulated by CaMKIV, Whereas Inhibitory Synaptic Scaling Is Under Independent Control. J Neurosci 37, 6778–6785.

Kaneko, M., Hanover, J.L., England, P.M., and Stryker, M.P. (2008a). TrkB kinase is required for recovery, but not loss, of cortical responses following monocular deprivation. Nat Neurosci 11, 497–504.

Kaneko, M., Stellwagen, D., Malenka, R.C., and Stryker, M.P. (2008b). Tumor necrosis factor-alpha mediates one component of competitive, experience-dependent plasticity in developing visual cortex. Neuron 58, 673–680.

Katz, L.C., and Shatz, C.J. (1996). Synaptic activity and the construction of cortical circuits. Science 274, 1133–1138.

Kenny, K., Royer, L., Moore, A.R., Chen, X., Marr, M.T.I., and Paradis, S. (2017). Rem2 promotes synapse formation by regulating the expression of genes encoding synaptogenic molecules. Mol Cell Neurosci. In Press.

Kirkwood, A., Rioult, M.G., and Bear, M.F. (1996). Experience-dependent modification of synaptic plasticity in visual cortex. Nature 381, 526–528.

Kuhlman, S.J., Olivas, N.D., Tring, E., Ikrar, T., Xu, X., and Trachtenberg, J.T. (2013). A disinhibitory microcircuit initiates critical-period plasticity in the visual cortex. Nature 501, 543–546.

Lambo, M.E., and Turrigiano, G.G. (2013). Synaptic and intrinsic homeostatic mechanisms cooperate to increase L2/3 pyramidal neuron excitability during a late phase of critical period plasticity. J Neurosci 33, 8810–8819.

Lehmann, K., and Lowel, S. (2008). Age-dependent ocular dominance plasticity in adult mice. PLoS One 3, e3120.

Liput, D.J., Lu, V.B., Davis, M.I., Puhl, H.L., and Ikeda, S.R. (2016). Rem2, a member of the RGK family of small GTPases, is enriched in nuclei of the basal ganglia. Sci Rep 6, 25137.

Liu, Z., Golowasch, J., Marder, E., and Abbott, L.F. (1998). A model neuron with activity-dependent conductances regulated by multiple calcium sensors. J Neurosci 18, 2309–2320.

Luscher, C., and Malenka, R.C. (2011). Drug-evoked synaptic plasticity in addiction: from molecular changes to circuit remodeling. Neuron 69, 650–663.

Maffei, A., and Turrigiano, G.G. (2008). Multiple modes of network homeostasis in visual cortical layer 2/3. J Neurosci 28, 4377–4384.

Marder, E., and Goaillard, J.M. (2006). Variability, compensation and homeostasis in neuron and network function. Nat Rev Neurosci 7, 563–574.

Marder, E., and Prinz, A.A. (2003). Current compensation in neuronal homeostasis. Neuron 37, 2–4.

Mardinly, A.R., Spiegel, I., Patrizi, A., Centofante, E., Bazinet, J.E., Tzeng, C.P., Mandel-Brehm, C., Harmin, D.A., Adesnik, H., Fagiolini, M., and Greenberg, M.E. (2016). Sensory experience regulates cortical inhibition by inducing IGF1 in VIP neurons. Nature 531, 371–375.

Martin, S.J., Grimwood, P.D., and Morris, R.G. (2000). Synaptic plasticity and memory: an evaluation of the hypothesis. Annu Rev Neurosci 23, 649–711.

Mazurek, M., Kager, M., and Van Hooser, S.D. (2014). Robust quantification of orientation selectivity and direction selectivity. Front Neural Circuits 8, 92.

McCurry, C.L., Shepherd, J.D., Tropea, D., Wang, K.H., Bear, M.F., and Sur, M. (2010). Loss of Arc renders the visual cortex impervious to the effects of sensory experience or deprivation. Nat Neurosci 13, 450–457.

Moore, A.R., Ghiretti, A.E., and Paradis, S. (2013). A loss-of-function analysis reveals that endogenous Rem2 promotes functional glutamatergic synapse formation and restricts dendritic complexity. PLoS One 8, e74751.

Mrsic-Flogel, T.D., Hofer, S.B., Ohki, K., Reid, R.C., Bonhoeffer, T., and Hubener, M. (2007). Homeostatic regulation of eye-specific responses in visual cortex during ocular dominance plasticity. Neuron 54, 961–972.

Nataraj, K., Le Roux, N., Nahmani, M., Lefort, S., and Turrigiano, G. (2010). Visual deprivation suppresses L5 pyramidal neuron excitability by preventing the induction of intrinsic plasticity. Neuron 68, 750–762.

Nedivi, E. (1999). Molecular analysis of developmental plasticity in neocortex. J Neurobiol 41, 135–147.

O'Leary, T., Williams, A.H., Franci, A., and Marder, E. (2014). Cell types, network homeostasis, and pathological compensation from a biologically plausible ion channel expression model. Neuron 82, 809–821.

Pang, C., Crump, S.M., Chang, L., Correll, R.N., Finlin, B.S., Satin, J., and Andres, D. (2010). Rem GTPase interacts with the proximal CaV1.2 C-terminus and modulates calcium-dependent channel inactivation. Channels (Austin) 4.

Picard, N., Leslie, J.H., Trowbridge, S.K., Subramanian, J., Nedivi, E., and Fagiolini, M. (2014). Aberrant development and plasticity of excitatory visual cortical networks in the absence of cpg15. J Neurosci 34, 3517–3522.

Piochon, C., Kano, M., and Hansel, C. (2016). LTD-like molecular pathways in developmental synaptic pruning. Nat Neurosci 19, 1299–1310.

Pozo, K., and Goda, Y. (2010). Unraveling mechanisms of homeostatic synaptic plasticity. Neuron 66, 337–351.

Pratt, K.G., and Aizenman, C.D. (2007). Homeostatic regulation of intrinsic excitability and synaptic transmission in a developing visual circuit. J Neurosci 27, 8268–8277.

Ranson, A., Cheetham, C.E., Fox, K., and Sengpiel, F. (2012). Homeostatic plasticity mechanisms are required for juvenile, but not adult, ocular dominance plasticity. Proc Natl Acad Sci U S A 109, 1311–1316.

Redmond, L., Kashani, A.H., and Ghosh, A. (2002). Calcium regulation of dendritic growth via CaM kinase IV and CREB-mediated transcription. Neuron 34, 999–1010.

Ringach, D.L., Shapley, R.M., and Hawken, M.J. (2002). Orientation selectivity in macaque V1: diversity and laminar dependence. J Neurosci 22, 5639–5651.

Rittenhouse, C.D., Shouval, H.Z., Paradiso, M.A., and Bear, M.F. (1999). Monocular deprivation induces homosynaptic long-term depression in visual cortex. Nature 397, 347–350.

Royer, L., Herzog, J.J., Kenny, K., Tzvetkova, B., Cochrane, J.C., Marr, M.T., and Paradis, S. (2017). The Ras-like GTPase Rem2 is a potent endogenous inhibitor of calcium/calmodulin-dependent kinase II activity. bioRxiv.

Rudy, B., Fishell, G., Lee, S., and Hjerling-Leffler, J. (2011). Three groups of interneurons account for nearly 100% of neocortical GABAergic neurons. Dev Neurobiol 71, 45–61.

Rutherford, L.C., Nelson, S.B., and Turrigiano, G.G. (1998). BDNF has opposite effects on the quantal amplitude of pyramidal neuron and interneuron excitatory synapses. Neuron 21, 521–530.

Sawtell, N.B., Frenkel, M.Y., Philpot, B.D., Nakazawa, K., Tonegawa, S., and Bear, M.F. (2003). NMDA receptor-dependent ocular dominance plasticity in adult visual cortex. Neuron 38, 977–985.

Shatz, C.J. (2009). MHC class I: an unexpected role in neuronal plasticity. Neuron 64, 40–45.

Shepherd, J.D., Rumbaugh, G., Wu, J., Chowdhury, S., Plath, N., Kuhl, D., Huganir, R.L., and Worley, P.F. (2006). Arc/Arg3.1 mediates homeostatic synaptic scaling of AMPA receptors. Neuron 52, 475–484.

Siegel, M., Marder, E., and Abbott, L.F. (1994). Activity-dependent current distributions in model neurons. Proc Natl Acad Sci U S A 91, 11308–11312.

Skarnes, W.C., Rosen, B., West, A.P., Koutsourakis, M., Bushell, W., Iyer, V., Mujica, A.O., Thomas, M., Harrow, J., Cox, T., et al. (2011). A conditional knockout resource for the genome-wide study of mouse gene function. Nature 474, 337–342.

Smith, G.B., Heynen, A.J., and Bear, M.F. (2009). Bidirectional synaptic mechanisms of ocular dominance plasticity in visual cortex. Philos Trans R Soc Lond B Biol Sci 364, 357–367.

Southwell, D.G., Froemke, R.C., Alvarez-Buylla, A., Stryker, M.P., and Gandhi, S.P. (2010). Cortical plasticity induced by inhibitory neuron transplantation. Science 327, 1145–1148.

Syken, J., Grandpre, T., Kanold, P.O., and Shatz, C.J. (2006). PirB restricts ocular-dominance plasticity in visual cortex. Science 313, 1795–1800.

Titley, H.K., Brunel, N., and Hansel, C. (2017). Toward a Neurocentric View of Learning. Neuron 95, 19–32.

Turrigiano, G. (2012). Homeostatic synaptic plasticity: local and global mechanisms for stabilizing neuronal function. Cold Spring Harb Perspect Biol 4, a005736.

Turrigiano, G.G. (2008). The self-tuning neuron: synaptic scaling of excitatory synapses. Cell 135, 422–435.

Turrigiano, G.G., Leslie, K.R., Desai, N.S., Rutherford, L.C., and Nelson, S.B. (1998). Activity-dependent scaling of quantal amplitude in neocortical neurons. Nature 391, 892–896.

Valverde, F. (1971). Rate and extent of recovery from dark rearing in the visual cortex of the mouse. Brain Res 33, 1–11.

van Versendaal, D., and Levelt, C.N. (2016). Inhibitory interneurons in visual cortical plasticity. Cell Mol Life Sci.

van Versendaal, D., Rajendran, R., Saiepour, M.H., Klooster, J., Smit-Rigter, L., Sommeijer, J.P., De-Zeeuw, C.I., Hofer, S.B., Heimel, J.A., and Levelt, C.N. (2012). Elimination of inhibitory synapses is a major component of adult ocular dominance plasticity. Neuron 74, 374–383.

Wallace, W., and Bear, M.F. (2004). A morphological correlate of synaptic scaling in visual cortex. J Neurosci 24, 6928–6938.

Yager, L.M., Garcia, A.F., Wunsch, A.M., and Ferguson, S.M. (2015). The ins and outs of the striatum: role in drug addiction. Neuroscience 301, 529–541.

Yoon, B.J., Smith, G.B., Heynen, A.J., Neve, R.L., and Bear, M.F. (2009). Essential role for a long-term depression mechanism in ocular dominance plasticity. Proc Natl Acad Sci U S A 106, 9860–9865.

